# HIV-1 Vpu restricts Fc-mediated effector functions in vivo

**DOI:** 10.1101/2022.02.21.481308

**Authors:** Jérémie Prévost, Sai Priya Anand, Jyothi Krishnaswamy Rajashekar, Jonathan Richard, Guillaume Goyette, Halima Medjahed, Gabrielle Gendron-Lepage, Hung-Ching Chen, Yaozong Chen, Joshua A. Horwitz, Michael W. Grunst, Susan Zolla-Pazner, Barton F. Haynes, Dennis R. Burton, Richard A. Flavell, Frank Kirchhoff, Beatrice H. Hahn, Amos B. Smith, Marzena Pazgier, Michel C. Nussenzweig, Priti Kumar, Andrés Finzi

## Abstract

Non-neutralizing antibodies (nnAbs) can eliminate HIV-1-infected cells via antibody-dependent cellular cytotoxicity (ADCC) and were identified as a correlate of protection in the RV144 vaccine trial. Fc-mediated effector functions of nnAbs were recently shown to alter the course of HIV-1 infection *in vivo* using a *vpu*-defective virus. Since Vpu is known to downregulate cell surface CD4, which triggers conformational changes in the viral envelope glycoprotein (Env), we ask whether the lack of Vpu expression was linked to the observed nnAbs activity. We found that restoring Vpu expression greatly reduces nnAb recognition of infected cells, rendering them resistant to ADCC responses. Moreover, administration of a nnAb in humanized mice reduces viral loads only in animals infected with a *vpu*-defective but not with a wildtype virus. Finally, nnAb Fc-effector functions are observed only on cells expressing Env in the “open” conformation. This work highlights the importance of Vpu-mediated evasion of humoral responses.

## INTRODUCTION

Human immunodeficiency virus type 1 (HIV-1) envelope glycoproteins (Env) mediate viral entry into target cells. Env is synthesized as a trimeric gp160 precursor, which is further cleaved into two subunits linked by non-covalent bonds: the exterior gp120 and the transmembrane gp41 subunits (Earl et al., 1990; Freed et al., 1989; McCune et al., 1988). During the entry process, the gp120 subunit sequentially interacts with the CD4 surface molecule as well as one of its co-receptors (CCR5 or CXCR4) (Kwong et al., 1998; Shaik et al., 2019; Wu et al., 2010a), allowing the gp41 subunit to mediate fusion between the viral and target cell lipid membranes (Chan et al., 1997; Weissenhorn et al., 1997). Since Env is the sole viral antigen exposed at the surface of virions and infected cells, it represents the main target for humoral responses. Fusion-competent Env trimers can sample different conformations. Broadly neutralizing antibodies (bNAbs) mainly recognize the pre-fusion “closed” conformation (Li et al., 2020; Lu et al., 2019; Munro et al., 2014; Stadtmueller et al., 2018), while non-neutralizing antibodies (nnAbs) mainly bind to “open” CD4-bound conformations (Alsahafi et al., 2019; Jette et al., 2021; Munro et al., 2014; Yang et al., 2019). Primary difficult-to-neutralize HIV-1 isolates favor the pre-fusion “closed” conformation, thus exposing Env regions that are heavily glycan shielded (Doores, 2015; Lee et al., 2016; Li et al., 2020).

Since antiretroviral therapy (ART) is unable to eradicate HIV-1 reservoirs, monotherapy or combination of bNAbs targeting the CD4-binding site (3BNC117, VRC01, VRC07-523), the V3 glycan supersite (10-1074, PGT121) and the V2 apex (PGDM1400) are currently under investigation in multiple clinical trials as therapeutic agents to reduce or eliminate cellular reservoirs through Fc-mediated effector functions (NCT02140255, NCT03837756, NCT04319367, NCT03721510). Thus far, results have shown that bNAbs can control HIV-1 viremia and delay viral rebound upon treatment interruption in HIV-1 infected individuals (Bar et al., 2016; Caskey et al., 2015; Caskey et al., 2017; Lynch et al., 2015; Scheid et al., 2016). Similar outcomes were also observed in non-human primates (NHP) and humanized mice (Barouch et al., 2013; Bolton et al., 2016; Freund et al., 2017; Halper-Stromberg et al., 2014; Hessell et al., 2016; Horwitz et al., 2013; Klein et al., 2012; Nishimura et al., 2017; Parsons et al., 2019; Schommers et al., 2020; Schoofs et al., 2019; Shingai et al., 2013). *In vivo* studies in animal models have demonstrated that Fc-mediated effector functions are required for the optimal therapeutic activity of bNAbs (Asokan et al., 2020; Bournazos et al., 2014; Halper-Stromberg et al., 2014; Lu et al., 2016; Wang et al., 2020). However, bNAbs rarely arise during natural infection and have yet to be consistently elicited by vaccination (Landais and Moore, 2018; Pauthner et al., 2019).

Given the difficulty of eliciting bNAbs *in vivo*, nnAbs have been evaluated as a potential alternative. nnAbs represent the majority of antibodies in the plasma of HIV-1-infected individuals and are easily elicited by vaccination (Beaudoin-Bussieres et al., 2020; Davis et al., 2009; Decker et al., 2005; Madani et al., 2016; Madani et al., 2018; Tomaras and Haynes, 2009; Tomaras et al., 2008; Visciano et al., 2019). Despite poor neutralization capacity, nnAbs can mediate other functions, such as the elimination of HIV-1-infected cells by antibody-dependent cellular cytotoxicity (ADCC) or antibody-dependent cellular phagocytosis (ADCP). Among these functions, ADCC was associated with the protection observed in the RV144 vaccine trial (Haynes et al., 2012a; Rerks-Ngarm et al., 2009). Thus, several studies have examined the antiviral effects of nnAbs in non-human primates and humanized mice. These included prophylactic administration of nnAbs targeting the CD4-binding site (b6), the V3 loop (KD-247, 2219), the V1V2 region (830A, 2158), the gp120 cluster A (A32) and the gp41 immunodominant region (246D, 4B3, F240, 7B2). The results from these studies showed a reduction in the number of transmitted/founder (T/F) viruses and/or plasma viral loads after challenge with lab-adapted tier 1 viruses (Burton et al., 2011; Eda et al., 2006; Hessell et al., 2018; Hioe et al., 2022; Moog et al., 2014; Santra et al., 2015). Finally, therapeutic administration of large quantities of the monoclonal anti-gp41 246D nnAb to humanized mice infected with a *vpu*-deleted tier 2 HIV-1 molecular clone (HIV-1_NL4/3_YU2) led to the elimination of infected cells and selected for escape mutations that stabilized Env “closed” conformation in an Fc-dependent manner, suggesting a protective effect (Horwitz et al., 2017). However, other studies administrating a cocktail of nnAbs (A32 and 17b) to humanized mice infected with a wild-type tier 2 HIV-1 strain (JR-CSF) had no impact on viral replication, except when combined with a small CD4 mimetic compound (CD4mc) that “open-up” the trimer and expose these otherwise occluded epitopes (Rajashekar et al., 2021).

Differences between the various nnAb studies could be attributed to the specificity of the antibodies (anti-gp41 vs anti-gp120), the humanized mouse model used or specific viral determinants. In particular, certain studies conducted using an infectious molecular clone (IMC) of HIV-1 that expressed a tier 2 Env (YU2) lacked a functional Vpu (HIV-1_NL4/3_YU2). However, Vpu plays an important role in downregulating cell surface CD4, which can bind and trigger conformational changes in the viral Env, therefore exposing vulnerable epitopes (Prevost et al., 2022; Veillette et al., 2015; Veillette et al., 2014). We thus asked whether the lack of a functional Vpu was responsible for the observed nnAbs activity by comparing isogenic viruses that differed solely in Vpu expression. Importantly, we tested the influence of Vpu expression on nnAb function not only *in vitro* but also in humanized mice. We found that anti-gp41 246D nnAb mediates potent ADCC responses against HIV-1_NL4/3_YU2, but not its Vpu+ counterpart. Accordingly, 246D was found to alter viral replication *in vivo* only in absence of Vpu. These data thus provide conclusive evidence that Vpu allows HIV-1 to evade humoral responses and emphasizes the need to use fully functional IMCs to assess nnAb Fc-mediated effector functions.

## RESULTS

### Elicitation of anti-gp41 non-neutralizing antibodies following immunization or HIV-1-infection

First, we characterized the susceptibility of the HIV-1_NL4/3_YU2 IMC to nnAbs-mediated Fc-effector responses to confirm previous observations (Horwitz et al., 2017). To do so, we used plasma samples from a cross-sectional cohort of 50 HIV-1-infected individuals (HIV+ plasma) which were grouped according to the inferred time post-infection and ART treatment (**Table S1**). The nnAbs present in the HIV+ plasma from this cohort were previously shown to mediate potent ADCC responses against infected cells presenting Env in the “open” CD4-bound conformation (Ding et al., 2016a; Richard et al., 2015; Veillette et al., 2015). We infected primary CD4+ T cells with the HIV-1_NL4/3_YU2 IMC and evaluated the ability of HIV+ plasma to recognize and eliminate infected cells. Consistent with its susceptibility to nnAbs, HIV-1_NL4/3_YU2-infected primary CD4+ T cells were efficiently recognized (**Figure 1A**) and susceptible to ADCC (**Figure 1B**) mediated by all the tested plasma samples, with a significant increase of activity starting six months post-infection. Similar levels of activity were also present in plasma from ART-treated individuals (**Figure 1A-B**).

**Figure 1.**
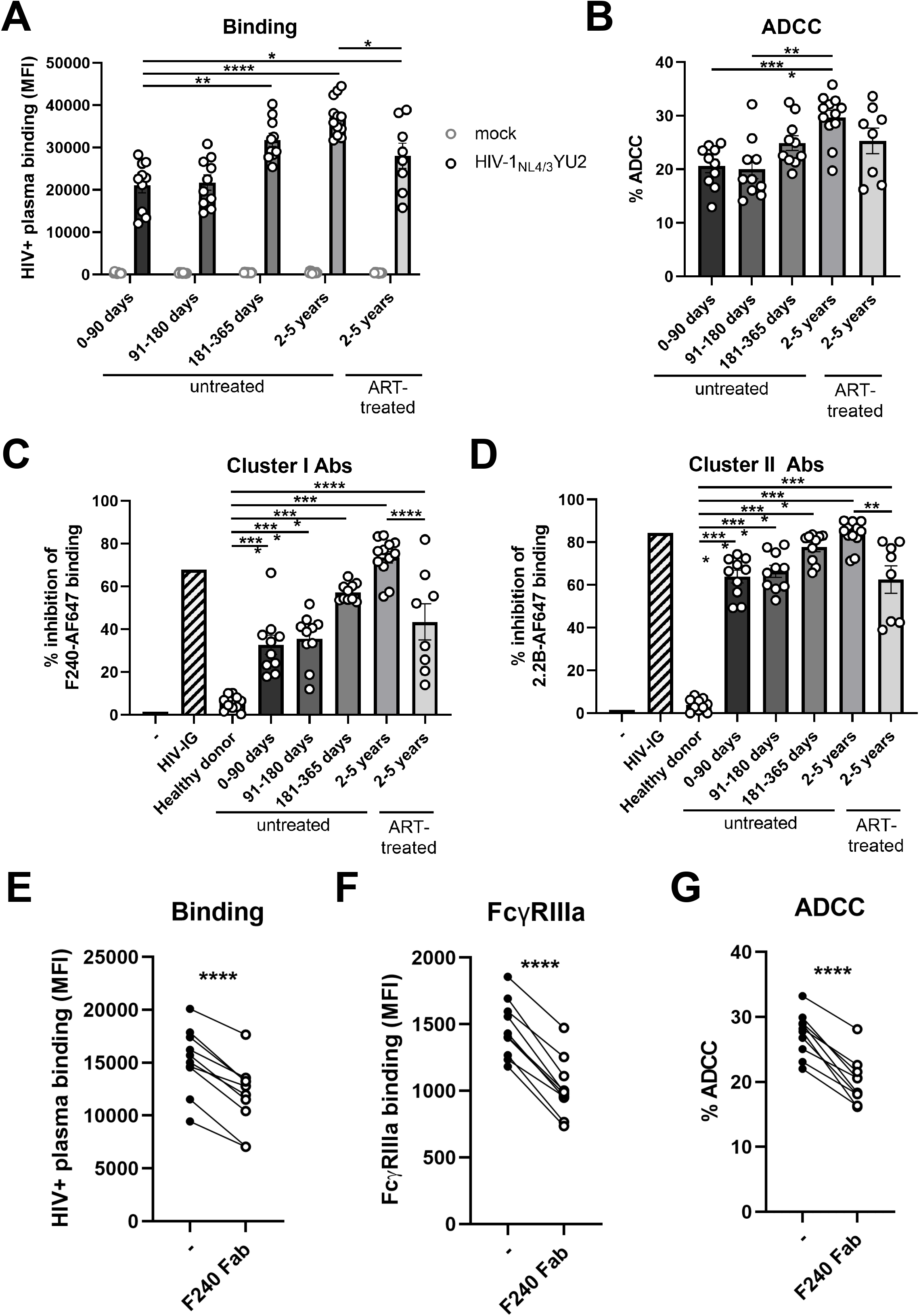
Cells infected with HIV-1_NL4/3_YU2 are susceptible to nnAb-mediated ADCC responses. Primary CD4+ T cells were either mock-infected (grey symbols) or infected with the HIV-1_NL4/3_YU2 virus (black symbols). (**A**) Cell-surface staining and (**B**) ADCC responses mediated by plasma from 50 different HIV-1-infected individuals categorized by the time post-infection and the antiretroviral therapy (ART) treatment status. (**A**) The graph shows the mean fluorescence intensities (MFI) on the whole cell population (mock) or infected p24+ cell population (HIV-1_NL4/3_YU2). (**C-D**) The binding of Alexa Fluor 647 (AF647)-precoupled (**C**) anti-gp41 cluster I F240 mAb or (**D**) anti-gp41 cluster II 2.2B mAb was evaluated in presence of unlabeled human plasma from healthy controls or HIV-1-infected individuals. A pool of purified immunoglobulins from HIV-1-infected individuals (HIV-IG) was used as a positive control. (**E**) Plasma binding, (**F**) FcγRIIIa binding and (**G**) ADCC responses mediated by plasma from 10 untreated HIV-1-infected individuals (2-5 years) were evaluated in presence of competing anti-gp41 cluster I F240 Fab fragment. Error bars indicate means ± standard errors of the means (SEM). Statistical significance was tested using (**A-D**) one-way ANOVA with a Holm-Sidak post-test or (**E-G**) a paired t test (*, P < 0.05; **, P < 0.01; ***, P < 0.001; ****, P < 0.0001; ns, nonsignificant).

Since the HIV-1_NL4/3_YU2 IMC has been reported to be sensitive to anti-gp41 Fc-mediated antiviral activity *in vivo* (Horwitz et al., 2017), we evaluated the contribution of anti-gp41 nnAbs present in HIV+ plasma to infected-cell recognition and ADCC activity by performing binding competition experiments (**Figure 1C-D**). We focused on two main classes of anti-gp41 nnAbs based on observations from previous studies showing potent ADCC responses (Ding et al., 2016a; Gohain et al., 2016; Moog et al., 2014; Santra et al., 2015; Sojar et al., 2019; von Bredow et al., 2016; Williams et al., 2019; Yang et al., 2018): anti-cluster I Abs targeting the disulfide loop region (C-C loop) (Gohain et al., 2016; Santra et al., 2015) and the anti-cluster II Abs targeting the heptad repeat region 2 (HR2) (Frey et al., 2010) (**Figure S1A**). Anti-gp41 cluster II nnAbs, inferred from binding competition experiments using the prototypic anti-cluster II 2.2B nnAb, were elicited in the acute phase of the infection (within 90 days) (**Figure 1D**). Elicitation of anti-gp41 cluster I nnAbs appears to take more time as revealed in binding competition experiments using the prototypic F240 anti-cluster I nnAb. While some competition with F240 binding was observed within the first three months post-infection, it culminates in the chronic phase of the infection (more than 2 years) (**Figure 1C**). In agreement with previous studies showing potent ADCC activity by anti-cluster I gp41 monoclonal Abs (Ding et al., 2016a; Gohain et al., 2016; Moog et al., 2014; Santra et al., 2015; von Bredow et al., 2016; Williams et al., 2019), we observed that blockade with anti-cluster I gp41 F240 Fab fragment significantly decreased plasma binding, FcγRIIIa engagement and ADCC responses against HIV-1_NL4/3_YU2 infected cells (**Figure 1E-G**).

### Vpu protects HIV-1-infected cells from recognition and Fc-effector functions mediated by anti-Env antibodies *in vitro* and *ex vivo*

Since the HIV-1_NL4/3_YU2 does not express the accessory protein Vpu due to a mutation in the start codon of the *vpu* gene (Horwitz et al., 2017), we asked whether the efficient recognition and ADCC-mediated elimination of HIV-1_NL4/3_YU2 infected primary CD4+ T cells by nnAbs was linked to the lack of Vpu expression by this IMC. We restored the *vpu* open reading frame (ORF, **Figure 2A**) and infected primary CD4+ T cells using both Vpu-negative (Vpu-) and Vpu-positive (Vpu+) constructs. Using a recently developed FACS-based intracellular staining method (Prevost et al., 2022), we confirmed Vpu expression upon restoration of the *vpu* ORF (**Figure 2B**). We also confirmed by intracellular staining the equivalent expression of Nef in both IMCs (**Figure 2B**). Vpu efficiently downregulated cell-surface CD4 and BST-2 (**Figure 2C**). We also measured cell surface expression of NTB-A and PVR, which were shown to modulate ADCC responses against HIV-1-infected cells (Prevost et al., 2019) and are downregulated by Vpu (Matusali et al., 2012; Shah et al., 2010). As expected, Vpu expression significantly downregulated their expression from the surface of primary CD4+ T cells from five different healthy individuals (**Figure 2D-E**).

**Figure 2.**
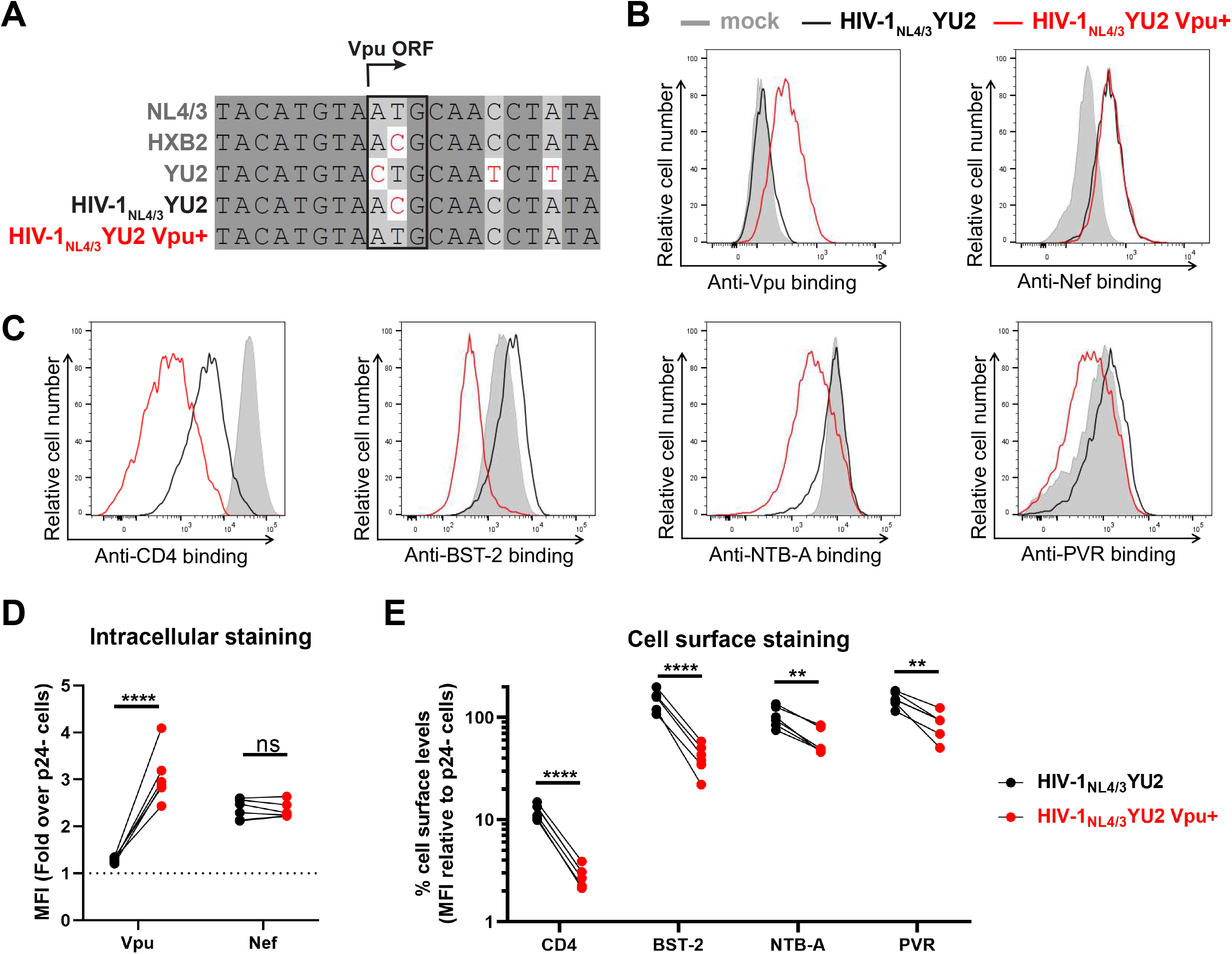
Reversion of Vpu open reading frame in the HIV-1_NL4/3_YU2 construct. (**A**) Sequence alignment of the Vpu open reading frame (ORF) region of selected HIV-1 reference isolates (NL4/3, HXB2, YU2) with the HIV-1_NL4/3_YU2 construct and its Vpu+ counterpart. Identical nucleotides are shaded in dark gray, conserved nucleotides in light gray and non-identical nucleotides are highlighted in red. The *vpu* start codon is highlighted by a black box. (**B-E**) Primary CD4+ T cells either mock-infected or infected with the HIV-1_NL4/3_YU2 virus or its Vpu+ counterpart were stained for intracellular detection of (**B**,**D**) viral proteins Vpu and Nef and cell-surface detection of (**C**,**E**) cellular proteins CD4, BST-2, NTB-A and PVR. (**B**-**C**) Histograms depicting representative stainings for (**B**) viral proteins and (**C**) their target substrates. (**D**-**E**) The graphs show the mean fluorescence intensities (MFI) obtained from five independent experiments using primary cells from five different healthy donors. (**D**) Viral protein levels were reported as a fold increase in the signal detected on infected p24+ cells compared to uninfected p24-cells. (**E**) Cellular protein levels were reported as a percentage of detection at the surface of infected p24+ cells compared to uninfected p24-cells. Error bars indicate means ± standard errors of the means (SEM). Statistical significance was tested using an unpaired t test or a Mann-Whitney U test based on statistical normality (**, P < 0.01; ****, P < 0.0001; ns, nonsignificant).

We next evaluated the effect of Vpu on the recognition and elimination of infected cells by nnAbs. Consistent with the observed downmodulation of CD4 and BST-2, Vpu expression strongly reduced the recognition of infected cells by monoclonal antibodies (mAb) targeting the anti-gp41 cluster I (246D), the gp41 cluster II (M785U3) and the gp120 cluster A (A32) (**Figure 3A**). We extended these results to a panel of 27 nnAbs which yielded the same results (**Figure 3B-C**). As a measure of this panel of nnAbs to mediate Fc-effector functions, we examined their ability to interact with a soluble dimeric FcγRIIIa protein. This recombinant protein is used as a surrogate of FcγR clustering which is required to trigger Fc-effector functions (Anand et al., 2019; Wines et al., 2017; Wines et al., 2016). Consistent with a significant reduction in the recognition of infected cells by nnAbs, we observed that Vpu expression diminished the ability of all mAbs to engage with FcγRIIIa (**Figure 3D-E**) and to mediate ADCC (**Figure 3F-G**). We noted that nnAbs targeting the gp41 were more potent at engaging soluble dimeric FcγRIIIa and mediating ADCC against cells infected with the *vpu*-defective IMC than the panel of anti-gp120 nnAbs tested (**Figure S2A-C**).

**Figure 3.**
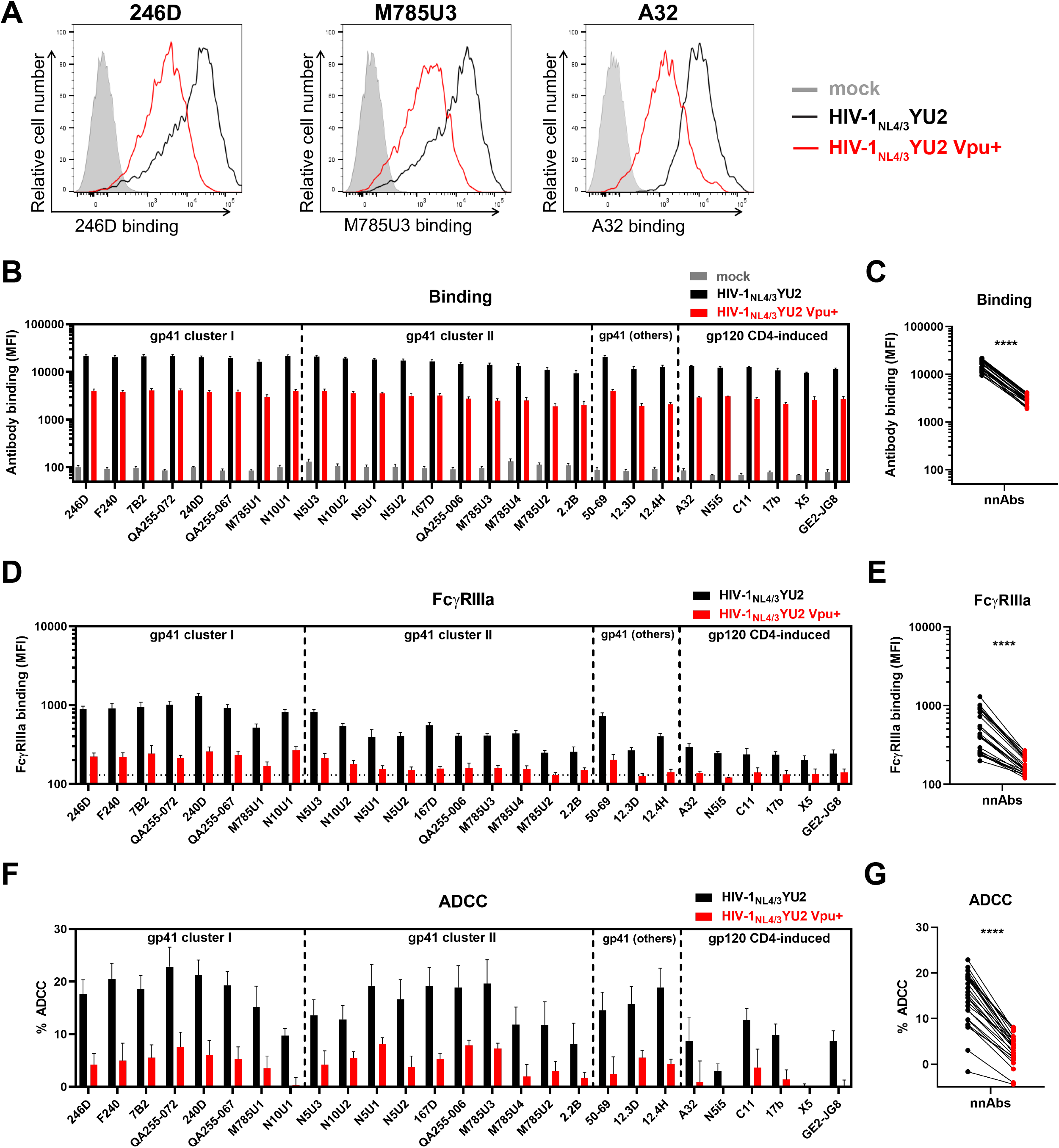
Vpu expression impairs Env recognition and Fc-effector functions mediated by anti-Env nnAbs. Primary CD4+ T cells were either mock-infected or infected with the HIV-1_NL4/3_YU2 virus or its Vpu+ counterpart. (**A-E**) Cell surface staining performed using anti-gp41 non-neutralizing antibodies targeting the gp41 cluster I, cluster II or other gp41 regions as well as anti-gp120 CD4-induced antibodies. Antibody binding was detected either by using (**A**-**C**) Alexa Fluor 647-conjugated anti-human secondary Abs or (**D**-**E**) by using biotinylated recombinant soluble dimeric FcγRIIIa followed by the addition of Alexa Fluor 647-conjugated streptavidin. (**A**) Histograms depicting representative stainings using anti-gp41 cluster I 246D, anti-gp41 cluster II M785U3 and anti-gp120 cluster A A32 mAbs. (**B**,**D**) The graphs represent the mean fluorescence intensities (MFI) obtained from the infected p24+ cell population using cells from five different healthy donors. (**D**) The horizontal dotted line represents the signal obtained in absence of mAb. (**F-G**) Primary CD4+ T cells infected with HIV-1_NL4/3_YU2 viruses were used as target cells. Autologous PBMCs were used as effector cells in a FACS-based ADCC assay. (**F**) The graph represents the percentages of ADCC obtained in the presence of the respective antibodies using cells from five different healthy donors. Error bars indicate means ± standard errors of the means (SEM). (**C**,**E,G**) Statistical significance was tested using a paired t test (****, P < 0.0001).

Vpu facilitates viral release by downregulating the restriction factor BST-2 (Neil et al., 2008; Van Damme et al., 2008). Multiple studies have shown that this activity decreases the overall amount of Env at the cell surface, consequently decreasing the susceptibility of infected cells to ADCC (Alvarez et al., 2014; Arias et al., 2014; Pham et al., 2016; Richard et al., 2017; Veillette ^e^t al., 201^4^). Since the *vpu*-defective HIV-1_NL4/3_YU2 was used to evaluate the *in vivo* activity of several bNAbs (Bournazos et al., 2016; Bournazos et al., 2014; Diskin et al., 2013; Freund et al., 2015; Freund et al., 2017; Halper-Stromberg et al., 2014; Horwitz et al., 2013; Klein et al., 2012; Klein et al., 2014; Lu et al., 2016; Schommers et al., 2020; Schoofs et al., 2019; Vanshylla et al., 2021), we tested whether their binding to infected cells was also influenced by Vpu expression. Consistent with decreased Env expression on cells infected with a Vpu+ virus, we observed a significant reduction in CD4-binding site (3BNC117), V3 glycan (10-1074) and V2-apex (PG16) bNAb binding (**Figure 4A**). The phenotype was validated using a panel of 35 bNAbs targeting different epitopes, indicative of an overall reduction of cell-surface Env expression in the presence of a functional Vpu (**Figure 4B-C**). Accordingly, infected cells were significantly less susceptible to bNAbs-mediated Fc-effector functions (**Figure 4D-G**). Among the different classes of bNAbs, antibodies targeting the V3 glycan supersite, the CD4-binding site and the V2 apex demonstrated stronger ADCC-mediated killing of cells infected with the *vpu*-defective HIV-1_NL4/3_YU2, compared with antibodies known to interact with the gp120 silent face, the gp120-gp41 interface or the gp41 membrane-proximal external region (MPER), despite similar levels of Env recognition (**Figure S2D-F**). These findings support previous observations indicating that in addition to Env recognition, the angle of approach of the antibody is important to mediate ADCC as it modulates the exposure of the Fc region required to activate effector cells (Acharya et al., 2014; Tolbert et al., 2020).

**Figure 4.**
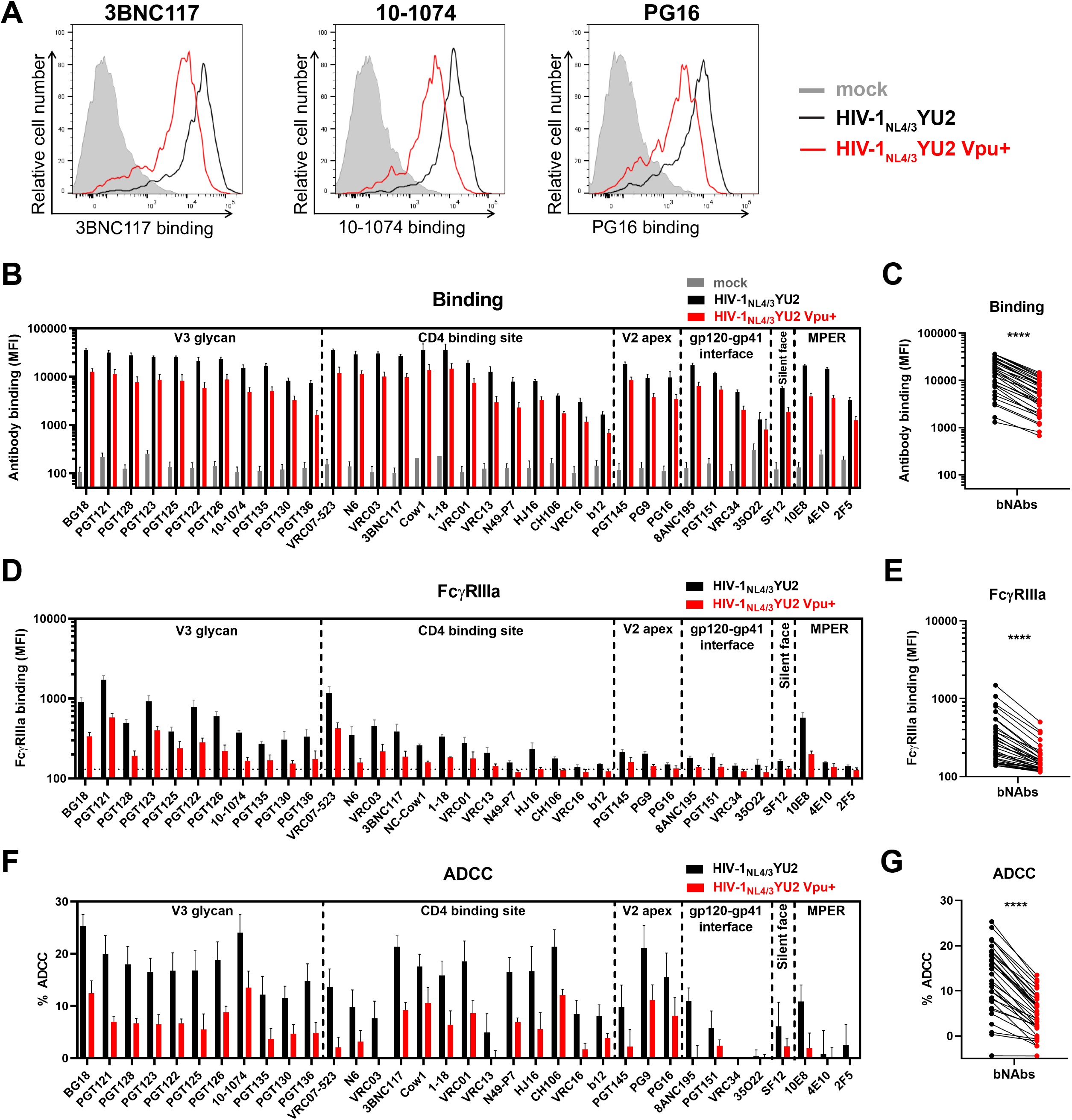
Vpu expression decreases Env recognition and Fc-effector functions mediated by anti-Env bNAbs. Primary CD4+ T cells were either mock-infected or infected with the HIV-1_NL4/3_YU2 virus or its Vpu+ counterpart. (**A-E**) Cell surface staining performed using broadly neutralizing antibodies targeting the gp120 V3 glycan supersite, CD4 binding site, V2 apex and silent face, the gp120-gp41 interface and the gp41 MPER. Antibody binding was detected either by using (**A**-**C**) Alexa Fluor 647-conjugated anti-human secondary Abs or (**D**-**E**) by using biotinylated recombinant soluble dimeric FcγRIIIa followed by the addition of Alexa Fluor 647-conjugated streptavidin. (**A**) Histograms depicting representative stainings using anti-CD4 binding site 3BNC117, anti-V3 glycan 10-1074 and anti-V2 apex PG16 mAbs. (**B**,**D**) The graphs represent the mean fluorescence intensities (MFI) obtained from the infected p24+ cell population using cells from five different healthy donors. (**D**) The horizontal dotted line represents the signal obtained in absence of mAb. (**F-G**) Primary CD4+ T cells infected with HIV-1_NL4/3_YU2 viruses were used as target cells. Autologous PBMCs were used as effector cells in a FACS-based ADCC assay. (**F**) The graph represents the percentages of ADCC obtained in the presence of the respective antibodies using cells from five different healthy donors. Error bars indicate means ± standard errors of the means (SEM). (**C**,**E,G**) Statistical significance was tested using a paired t test (****, P < 0.0001).

Since all coding regions of the HIV-1_NL4/3_YU2 IMC other than the *env* gene are derived from a lab-adapted proviral backbone, we wished to test unmodified HIV-1 primary isolates. Primary CD4+ T cells were infected with a panel of 13 infectious molecular clones coding for Vpu or not. This panel includes transmitted/founder (T/F) viruses, molecular clones derived during chronic infection, and IMCs from lab-adapted strains as controls (**Figure 5A-B**). Vpu expression consistently reduced nnAbs (246D and M785U3) and bNAbs (3BNC117 and 10-1074) recognition of infected cells, which in turn protected infected cells from ADCC responses mediated by all antibodies tested (**Figure 5A-B**). Env polymorphisms present in CH167 (E460), REJO (N334) and CH293 (T332, N334) viruses abrogated the capacity of bNAbs 3BNC117 or 10-1074 to recognize infected cells. Of note, while Vpu expression has a profound effect in the recognition of infected cells by all tested anti-Env antibodies, it did not affect their neutralization profile (**Figure S1B-C**).

**Figure 5.**
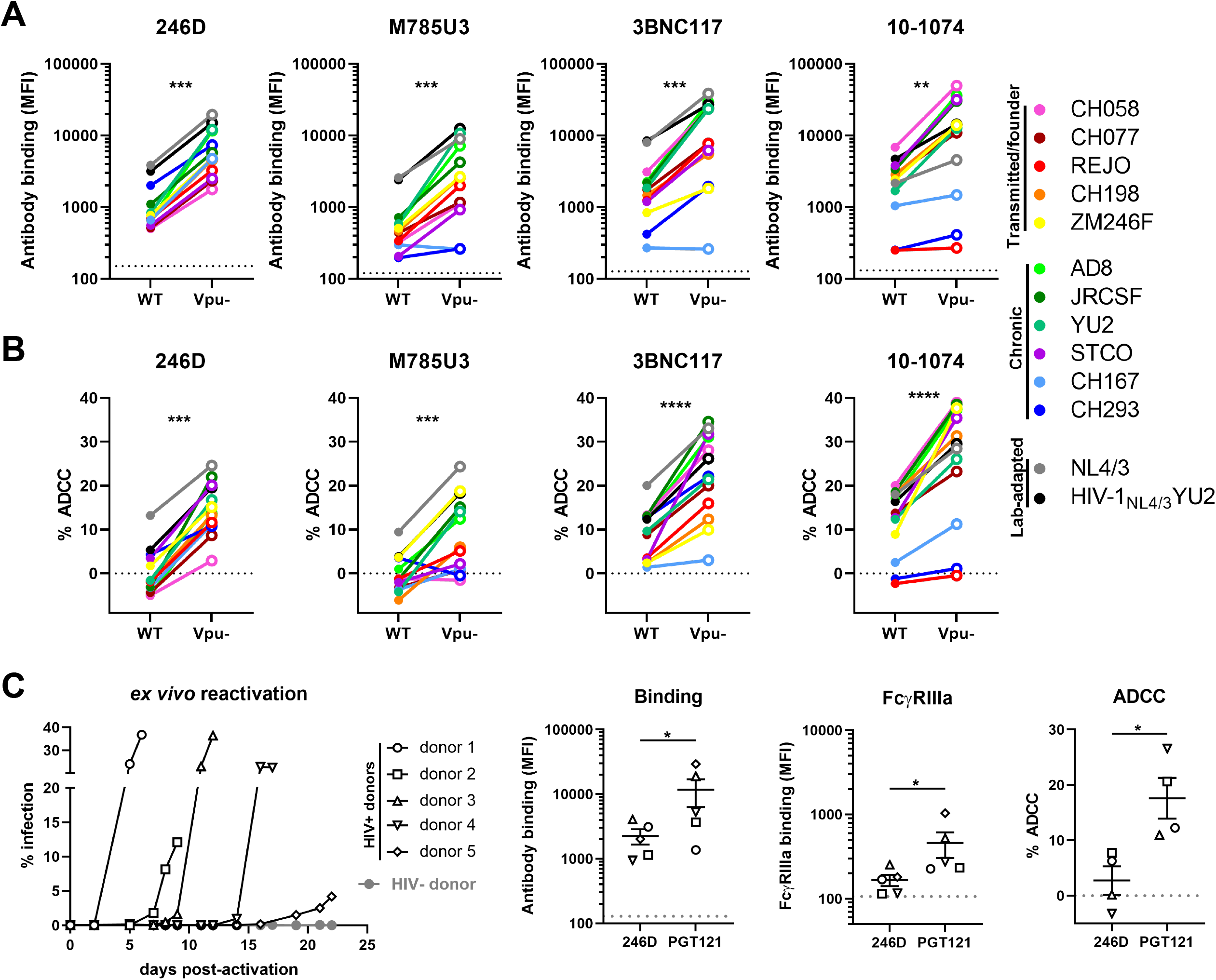
The ability of Vpu to limit anti-Env ADCC responses is conserved among different HIV-1 strains. (**A-B**) Primary CD4+ T cells were infected with HIV-1 clade B and clade C transmitted/founder (CH058, CH077, REJO, CH198, ZM246F), chronic (AD8, JRCSF, YU2, STCO, CH167, CH293) and lab-adapted (NL4/3, HIV-1_NL4/3_YU2) wild-type (WT) strains or their Vpu-counterpart. (**A**) Cell surface staining performed using anti-gp41 nnAbs 246D and M785U3, as well as bNAbs 3BNC117 and 10-1074. Antibody binding was detected either by using Alexa Fluor 647-conjugated anti-human secondary Abs. (**B**) Infected primary CD4+ T cells were used as target cells and autologous PBMCs were used as effector cells in a FACS-based ADCC assay. (**C**) Primary CD4+ T cells from five different ART-treated HIV-1-infected individuals were isolated and activated with PHA-L/IL-2 to expand the endogenous virus. Cell surface staining and ADCC experiments were performed upon reactivation. Antibody binding was detected using Alexa Fluor 647-conjugated anti-human secondary Abs or biotinylated recombinant soluble dimeric FcγRIIIa followed by the addition of Alexa Fluor 647-conjugated streptavidin. *Ex vivo*-expanded infected primary CD4+ T cells from four HIV-1-infected individuals were used as target cells and autologous PBMCs were used as effector cells in a FACS-based ADCC assay. ADCC susceptibility was only measured when the percentage of infection (p24+ cells) was higher than 10%. (**A,C**) The horizontal dotted lines represent the signal obtained in absence of mAb. (**A-C**) The antibody binding and FcγRIIIa graphs represent the mean fluorescence intensities (MFI) obtained from the infected p24+ cell population. The ADCC graphs represent the percentages of ADCC obtained in the presence of the respective antibodies. Error bars indicate means ± standard errors of the means (SEM). Statistical significance was tested using (**A-B**) a paired t test or Wilcoxon signed-rank test based on statistical normality or (**C**) a Mann-Whitney U test (*, P < 0.05; **, P < 0.01; ***, P < 0.001; ****, P < 0.0001; ns, nonsignificant).

To extend our observations to a more physiological model, we expanded patient-derived infected CD4+ T cells. Briefly, we isolated and activated primary CD4+ T cells from five ART-treated HIV-1-infected individuals. Viral replication upon reactivation was monitored by intracellular p24 staining and flow cytometry (**Figure 5C**). These endogenously-infected cells were protected from Fc-mediated effector functions mediated by nnAb 246D but remained susceptible to bNAb PGT121 (**Figure 5C**), in agreement with our *in vitro* results generated using a Vpu+ IMC (**Figure 5A-B**).

### Vpu expression limits the antiviral activity of the 246D antibody in HIV-1-infected humanized mice

To determine the impact of Vpu expression on 246D antiviral activities *in vivo*, we infected SRG-15 (SIRPA^h/m^ Rag2^-/-^ Il2rg^-/-^ IL15^h/m^) humanized mice (hu-mice) with HIV-1_NL4/3_YU2 or its Vpu-competent counterpart. Similar to a previously used humanized mouse model (Horwitz et al., 2017), this hu-mice model supports HIV-1 replication and Fc-effector functions *in vivo* (Herndler-Brandstetter et al., 2017; Rajashekar et al., 2021). Immunodeficient mice engrafted with human peripheral blood lymphocytes (hu-PBL) were infected intraperitoneally (I.P.) with 30,000 plaque-forming units (PFU) of either virus (**Figure 6A**). Half of each cohort received subcutaneous (S.C.) injections of the 246D nnAb at days 2, 4 and 6 post-infection. Infected humanized mice were then monitored for plasma viral loads (PVLs). Both viruses replicated efficiently in SRG-15 hu-PBL mice, reaching on average 1×10^7^ viral RNA copies/mL of plasma at 3 days post-infection (P.I.) (**Figure 6B**). While PVLs in mice infected with HIV-1_NL4/3_YU2 stabilized after day 3, infection with the Vpu+ variant further increased PVLs to 3.5×10^7^ copies/mL by day 10. A single administration of the 246D nnAb at day 2 resulted in a significant reduction in PVLs (∼36-fold decrease) in hu-mice infected with the *vpu*-defective virus but not significantly in those infected with the *vpu*-competent virus (**Figure 6B**). 246D maximal inhibitory effect (∼85-fold reduction) was reached 10 days post-infection with the *vpu*-defective virus. At the end of the experiment (day 11), 246D treatment induced on average a 41-fold decrease in PVLs in mice infected with the *vpu*-defective virus but no difference in hu-mice infected with the Vpu+ virus. These results highlight the role played by Vpu in promoting viral replication in presence of nnAbs.

**Figure 6.**
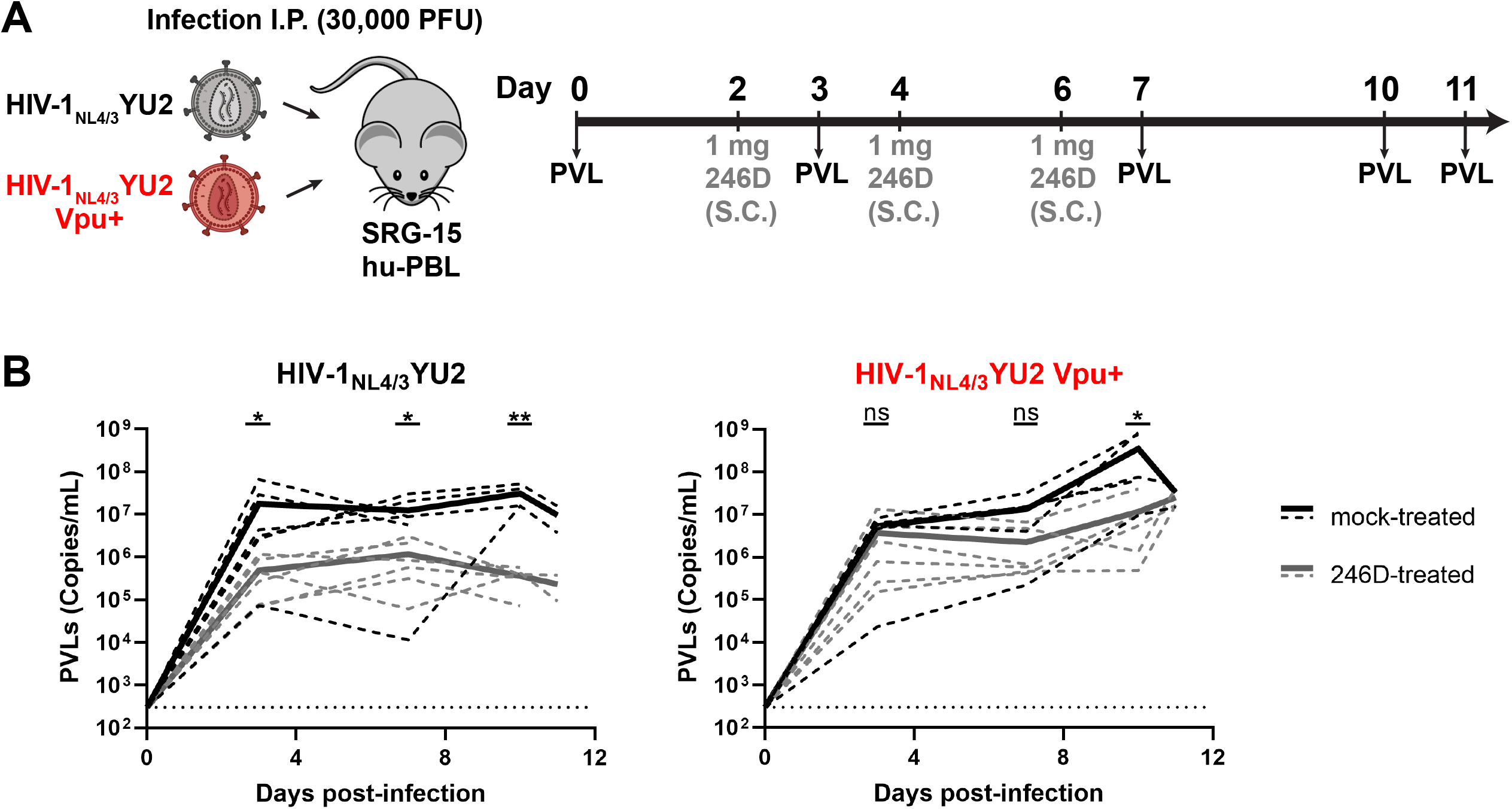
Vpu promotes HIV-1 replication in humanized mice treated with non-neutralizing antibody 246D. (**A**) Experimental outline. SRG-15-Hu-PBL mice were infected with HIV-1_NL4/3_YU2 or its Vpu+ counterpart intraperitoneally. At day 2, 4 and 6 post-infection, mice were administered 1mg of 246D mAb subcutaneously (S.C.). Mice were bled routinely for plasma viral load (PVL) quantification. (**B**) PVL levels were measured by quantitative real-time PCR (limit of detection = 300 copies/mL, dotted line). Twelve mice were infected using each virus; six of them were mock-treated (black lines) and the other six were treated with 246D mAb (gray lines). PVL measurements for individual mice are shown as dashed lines and mean values for each regimen are shown as solid lines. Statistical significance was tested using an unpaired t test or a Mann-Whitney U test based on statistical normality (*, P < 0.05;**, P < 0.01; ns, nonsignificant).

### Harnessing non-neutralizing antibody Fc-effector functions by improving epitope exposure and FcγRIIIa interaction

Enhancing the affinity of the Fc fragment of antibodies for FcγRs was shown to increase Fc-effector functions of bNAbs in HIV-1-infected humanized mice (Bournazos et al., 2014; Wang et al., 2020). To evaluate if this strategy could apply to nnAbs, we introduced well-characterized IgG1 Fc mutations in the 246D heavy chain to modulate its interaction with FcγRs. The GRLR mutations (G236R/L328R) and the GASDALIE mutations (G236A/S239D/A330L/I332E) are respectively known to decrease and increase the affinity for activating FcγRs (Bournazos et al., 2014; Horton et al., 2010; Smith et al., 2012). To characterize these Fc variants, primary CD4+ T cells were infected with the HIV-1_NL4/3_YU2 constructs expressing Vpu or not. As expected, Fc modifications did not alter the ability of 246D to recognize infected cells, but it modulated the interaction with the soluble dimeric FcγR probe, with the GRLR mutations abrogating FcγRIIIa binding and the GASDALIE mutations improving it (**Figure 7A-B**). Introduction of the GASDALIE mutations enhanced ADCC against cells infected with the *vpu*-defective virus. Interestingly, it also allowed 246D to mediate ADCC against cells infected with the Vpu+ IMC, while the unaltered native 246D failed to do so (**Figure 7C**). To evaluate if the 246D GASDALIE was able to mediate ADCC against cells infected with a primary isolate, we infected primary CD4+ T cells with the transmitted/founder virus CH058, a strain shown to be resistant to 246D-mediated ADCC responses (**Figure 5B**). CH058-infected cells were poorly recognized by 246D and resistant to ADCC mediated by all 246D Fc variants tested, including 246D GASDALIE (**Figure 7D-F**). Consistent with the role of Nef and Vpu in preventing the exposure of epitopes recognized by nnAbs, disruption of the expression of both accessory proteins enhanced recognition and ADCC susceptibility of infected cells by 246D (**Figure 7D-F**). These results agree with the requirement of Env-CD4 interaction to expose the gp41 cluster I region. 246D recognizes with picomolar affinity a highly conserved linear peptide (^596^WGCSGKLICTT^606^) which corresponds to the gp41 C-C loop, located between HR1 and HR2 helices (**Figure S3A-C**). This epitope is occluded in the “closed” Env trimer structure but can be exposed upon CD4-triggered conformational changes (**Figure S3D-E**).

**Figure 7.**
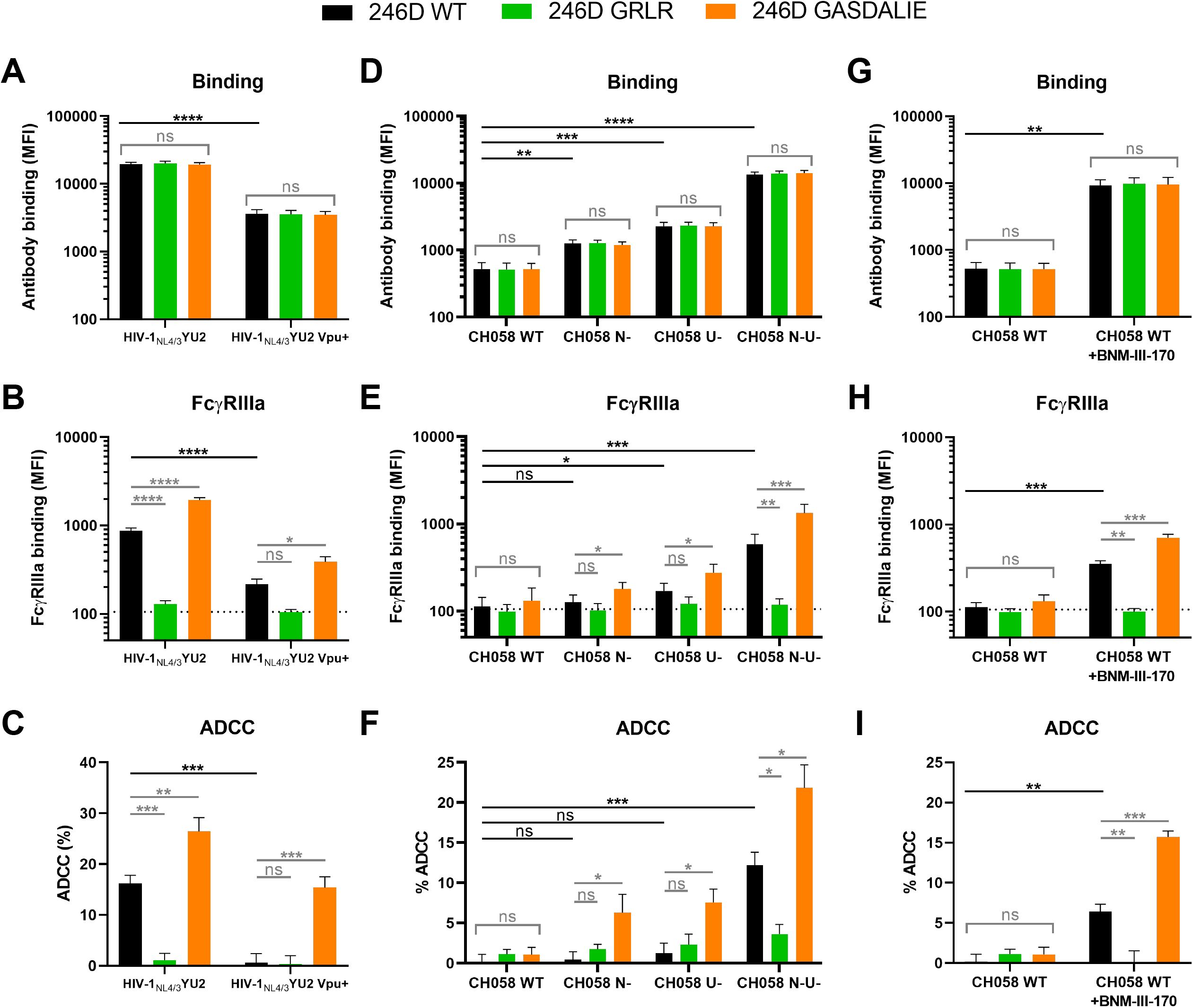
CD4 mimetics and Fc modifications boost the capacity of anti-gp41 nnAbs to mediate ADCC responses. Primary CD4+ T cells were either infected with (**A-C**) HIV-1_NL4/3_YU2 virus or its Vpu+ counterpart, (**D-F**) transmitted/founder virus CH058 wild-type (WT) or its Nef and/or Vpu-deleted counterparts (N-, U-, N-U-) or (**G**-**I**) CH058 WT with further treatment or not with CD4mc BNM-III-170 during staining and ADCC experiments. Cell surface staining performed using anti-gp41 nnAb 246D WT or Fc-mutated to impair (G236R/L328R; GRLR) or enhance (G236A/S239D/A330L/I332E; GASDALIE) Fc-effector functions. Antibody binding was detected using (**A**,**D**,**G**) Alexa Fluor 647-conjugated anti-human secondary Abs or (**B**,**E**,**H**) by using biotinylated recombinant soluble dimeric FcγRIIIa followed by the addition of Alexa Fluor 647-conjugated streptavidin. (**A**-**B**,**D**-**E**,**G-H**) The graphs represent the mean fluorescence intensities (MFI) obtained from the infected p24+ cell population using cells from five different healthy donors. (**B,E,H**) The horizontal dotted lines represent the signal obtained in absence of mAb. (**C**,**F**,**I**) Infected primary CD4+ T cells were used as target cells and autologous PBMCs were used as effector cells in a FACS-based ADCC assay. The graphs represent the percentages of ADCC obtained in the presence of the respective antibodies using cells from five different healthy donors. Error bars indicate means ± standard errors of the means (SEM). (**A-I**) Statistical significance was tested using a one-way ANOVA with a Holm-Sidak post-test or a Kruskal-Wallis test with a Dunn’s post-test when comparing between the different 246D Fc variants and (**C**) an unpaired t test or a Mann-Whitney U test when comparing between viruses or treatment. Appropriate statistical test (parametric or nonparametric) was applied based on dataset distribution normality (*, P < 0.05; **, P < 0.01; ***, P < 0.001; ****, P < 0.0001; ns, nonsignificant).

While it is becoming increasingly clear that HIV-1 successfully evades nnAbs responses by keeping its Env in a “closed” conformation (Bruel et al., 2017; Dufloo et al., 2020; Veillette et al., 2015; Veillette et al., 2014; von Bredow et al., 2016), new strategies are currently being tested to harness their potential antiviral activity. Small CD4 mimetic compounds have been optimized to “open up” Env trimers, therefore exposing otherwise occluded epitopes recognized by nnAbs (Ding et al., 2019b; Fritschi et al., 2021; Jette et al., 2021; Laumaea et al., 2020; Melillo et al., 2016). Using this strategy, CD4mc were shown to synergize with monoclonal CD4i Abs or nnAbs found in plasma from infected individuals to eliminate HIV-1-infected cells *in vitro*, *ex vivo* and *in vivo* in humanized mice (Alsahafi et al., 2019; Anand et al., 2019; Ding et al., 2016b; Lee et al., 2015; Madani et al., 2018; Prevost et al., 2020b; Rajashekar et al., 2021; Richard et al., 2016; Richard et al., 2017; Richard et al., 2015). Since 246D was unable to efficiently recognize cells infected with a primary isolate, we combined it with the CD4mc BNM-III-170, which greatly improved its capacity to bind to CH058-infected cells (**Figure 7G**). Accordingly, FcγRIIIa engagement and ADCC responses against WT-infected cells were significantly enhanced upon CD4mc addition (**Figure 7H-I**). Altogether, our results suggest that Fc-effector functions mediated by nnAbs are limited by the occluded nature of their epitopes.

## DISCUSSION

The HIV-1 accessory protein Vpu is a multi-functional protein that promotes viral replication by interfering with the intracellular trafficking of various host proteins (Dube et al., 2010). CD4 and BST-2 downregulation by Vpu was shown to increase viral release in cell culture systems (Bour et al., 1999; Neil et al., 2008; Van Damme et al., 2008). Humanized mouse models of acute HIV-1 infection using HIV-1-infected humanized mice confirmed the role of Vpu in promoting viral replication in the initial phase of infection (Dave et al., 2013; Sato et al., 2012; Yamada et al., 2018). Similarly, chronic infection of NHPs with SHIV constructs encoding a defective or mutated *vpu* gene were found to be less pathogenic and exhibit lower viral loads (Hout et al., 2005; Shingai et al., 2011). In these *in vivo* studies, elevated viral loads were linked to Vpu-mediated CD4 and BST-2 downregulation, but the potential contribution of nnAbs to PVLs were not addressed. Beyond its effect on viral release, Vpu protects HIV-1-infected cells from nnAbs-mediated ADCC responses by limiting the presence of Env-CD4 complexes at the plasma membrane (Prevost et al., 2022; Veillette et al., 2015; Veillette et al., 2014). Here, we show that Vpu enhances viral replication *in vivo* by limiting nnAbs recognition of infected cells and therefore their capacity to mediate Fc-effector functions. Beyond infected cell elimination, the absence of Vpu could also affect the level of circulating infectious viral particles. While we did not observe any changes in the neutralization by anti-gp41 nnAbs against virions produced in 293T cells *in vitro* (**Figure S1B-C**), one could speculate that the capacity of nnAbs to neutralize viral particles originating from primary cells could also be altered in the absence of Vpu. Indeed, CD4 incorporation into virions has been shown to sensitize them to neutralization by various CD4i mAbs and HIV-IG (Ding et al., 2019a). While this was observed with *nef*-defective viruses, CD4 incorporation is also modulated by Vpu expression and this could apply to *vpu*-defective viruses (Levesque et al., 2003). Our results highlight the importance of carefully selecting viruses for *in vitro* and *in vivo* studies, since several widely used HIV-1 strains are defective for Vpu expression (e.g. HXB2, YU2, ADA) (Li et al., 1991; Shaw et al., 1984; Theodore et al., 1996).

Different approaches aimed at generating bNAbs by vaccination are being investigated using germline-targeting Env immunogens (Dosenovic et al., 2015; Jardine et al., 2016; Steichen et al., 2016) followed by sequential Env trimer immunization (Escolano et al., 2016; Haynes et al., 2012b; Saunders et al., 2019; Williams et al., 2017) to guide antibody maturation. However, none of these vaccine studies have yet resulted in the consistent elicitation of bNAbs. Facing the difficulty in eliciting bNAbs by vaccination, nnAbs have been studied as an alternative for vaccine development. If ADCC activity contributes to vaccine protection, as suggested in the RV144 trial (Haynes et al., 2012a), our results suggest that elicitation of nnAbs are unlikely to confer protection in a vaccine setting unless strategies to “open” the Env trimer are in place. Supporting this, NHP immunized with monomeric gp120 were completely protected from a heterologous SHIV infection if a CD4mc was combined with the challenge viral stock (Madani et al., 2018). This study showed that nnAbs elicited by the gp120 immunogen did not protect in the absence of CD4mc.

In this study, we selected the SRG-15 humanized mouse model over other humanized mouse models because of its endogenous expression of human IL-15, which allows the development of a functional NK cell compartment (Herndler-Brandstetter et al., 2017). NK cells play a central role in the ADCC responses *in vitro* and *in vivo*, and their specific depletion has been shown to abrogate the elimination of HIV-1 reservoirs by a combination of CD4mc and nnAbs in SRG-15 humanized mice (Rajashekar et al., 2021). Human IL-15 expression likely also increases HIV-1 replication by regulating the susceptibility of CD4+ T cells to infection (Manganaro et al., 2018). This could explain the higher viral load peak (∼10^7^ copies/mL) (**Figure 6B**) compared to those achieved in the NRG humanized mice using the same HIV-1_NL4/3_YU2 IMC (∼10^5^ copies/mL) in previous studies (Halper-Stromberg et al., 2014; Horwitz et al., 2017). We also used humanized mice as a model of HIV-1 infection since it allows the use of clinically-relevant HIV-1 isolates and does not require Env adaptation to replicate.

In addition to accessory proteins, Env conformation is a critical factor when studying nnAbs Fc-effector functions (Alsahafi et al., 2016; Ding et al., 2016a; Dufloo et al., 2020; Prevost et al., 2021; Prevost et al., 2018a; Prevost et al., 2018b; Prevost et al., 2017; Richard et al., 2015; Veillette et al., 2015; Veillette et al., 2014). Multiple studies reported significant effects of nnAbs in limiting HIV-1/SHIV replication *in vivo* using viruses coding for easy-to-neutralize tier 1 Env from lab-adapted strains, which are readily recognized by nnAbs (Burton et al., 2011; Eda et al., 2006; Hessell et al., 2018; Hioe et al., 2022; Moog et al., 2014; Santra et al., 2015). A recent study showed that tier 2 primary virus can be impacted *in vivo* by nnAbs but only when combined with CD4mc (Rajashekar et al., 2021). This combination of nnAbs and CD4mc significantly reduced plasma viral loads and the size of the viral reservoir in an Fc-effector functions and NK cells dependent manner (Rajashekar et al., 2021). These results emphasize the need to utilize fully functional viruses in ADCC assays to preclude Vpu-related artifacts. However, one could speculate that the development of broad-spectrum Vpu inhibitors may enhance the efficacy of nnAbs to eliminate HIV-1-infected cells (Robinson et al., 2021). Overall, our study indicate that it is unlikely for nnAbs-based immunotherapies to alter viral replication in the absence of strategies aimed at exposing the vulnerable epitopes they recognize.

## ACKNOWLEDGMENTS

The authors thank the CRCHUM BSL3 and Flow Cytometry Platforms for technical assistance, and Mario Legault from the FRQS AIDS and Infectious Diseases network for cohort coordination and clinical samples. We thank the following collaborators for kindly providing antibodies: Julie Overbaugh (Fred Hutchinson Cancer Research Center) for QA255-006, QA255-067 and QA255-072, James Robinson (Tulane University) for 7B2, 2.2B, 12.3D, 12.4H, A32, C11 and 17b, George Lewis (University of Maryland) for M785U1, M785U2, M785U3, M785U4, N5U1, N5U2, N5U3, N10U1 and N10U2, Gunilla Karlsson Hedestam (Karolinska Institutet) for GE2-JG8, John Mascola (Vaccine Research Center, NIAID) for VRC01, VRC03, VRC07-523, VRC13, VRC16 and VRC34, Mark Connors (NIAID) for 10E8, N6 and 35O22, Florian Klein (University of Cologne) for 1-18, and the International AIDS Vaccine Initiative (IAVI) for PG9, PG16, PGT121, PGT122, PGT123, PGT125, PGT126, PGT128, PGT130, PGT135, PGT136, PGT145 and PGT151. We thank P. Mark Hogarth (Burnet Institute) for kindly providing recombinant dimeric FcγRIIIa. The Figure 6 was prepared using illustrations from <u>BioRender.com</u>.

This study was supported by a Canadian Institutes of Health Research (CIHR) foundation grant #352417 to A.F. Funds were also provided by a CIHR Team grant #422148 to P.K. and A.F., a Canada Foundation for Innovation (CFI) grant #41027 to A.F and by the National Institutes of Health to A.F. (R01 AI148379 and R01 AI150322), to M.P. and A.F. (R01 AI129769), M.P. (AI116274) and to P.K. (R01 AI145164, R33 AI122384 and P50 AI150464 [CHEETAH]). Support for this work was also provided by P01 GM56550/AI150471 to A.B.S. and A.F. This work was partially supported by 1UM1AI164562-01, co-funded by National Heart, Lung and Blood Institute, National Institute of Diabetes and Digestive and Kidney Diseases, National Institute of Neurological Disorders and Stroke, National Institute on Drug Abuse and the National Institute of Allergy and Infectious Diseases to A.F. A.F. is the recipient of a Canada Research Chair on Retroviral Entry #RCHS0235 950-232424. F.K. is supported by the German Research Foundation (DFG CRC 1279 and SPP 1923) and the Baden-Württemberg Foundation (BWST-ISF2018-032). J.P. and S.P.A. are recipients of CIHR doctoral fellowships. M.W.G. is a recipient of the Gruber Science Fellowship. The funders had no role in study design, data collection and analysis, decision to publish, or preparation of the manuscript.

## AUTHOR CONTRIBUTIONS

**Conceptualization:** J.P. and A.F.; **Methodology:** J.P., J.K.R., P.K. and A.F.; **Investigation:** J.P., S.P.A., J.K.R., J.R., G.G., H.M., G.G.-L., Y.C. and M.W.G.; **Resources:** J.P., H.-C.C., J.A.H., S.Z.-P., B.F.H., D.R.B., R.A.F., F.K., B.H.H., A.B.S., M.P., M.C.N., P.K. and A.F.; **Formal Analysis:** J.P.; **Visualization:** J.P.; **Supervision:** A.B.S., M.P., M.C.N., P.K. and A.F.; **Funding acquisition:** A.B.S., M.P., P.K. and A.F.; **Writing – original draft:** J.P., B.H.H. and A.F.; **Writing – review & editing:** All authors.

## DISCLAIMER

The opinions and assertions expressed herein are those of the author(s) and do not necessarily reflect the official policy or position of the Uniformed Services University or the Department of Defense.

## STAR METHODS

### KEY RESOURCES TABLE

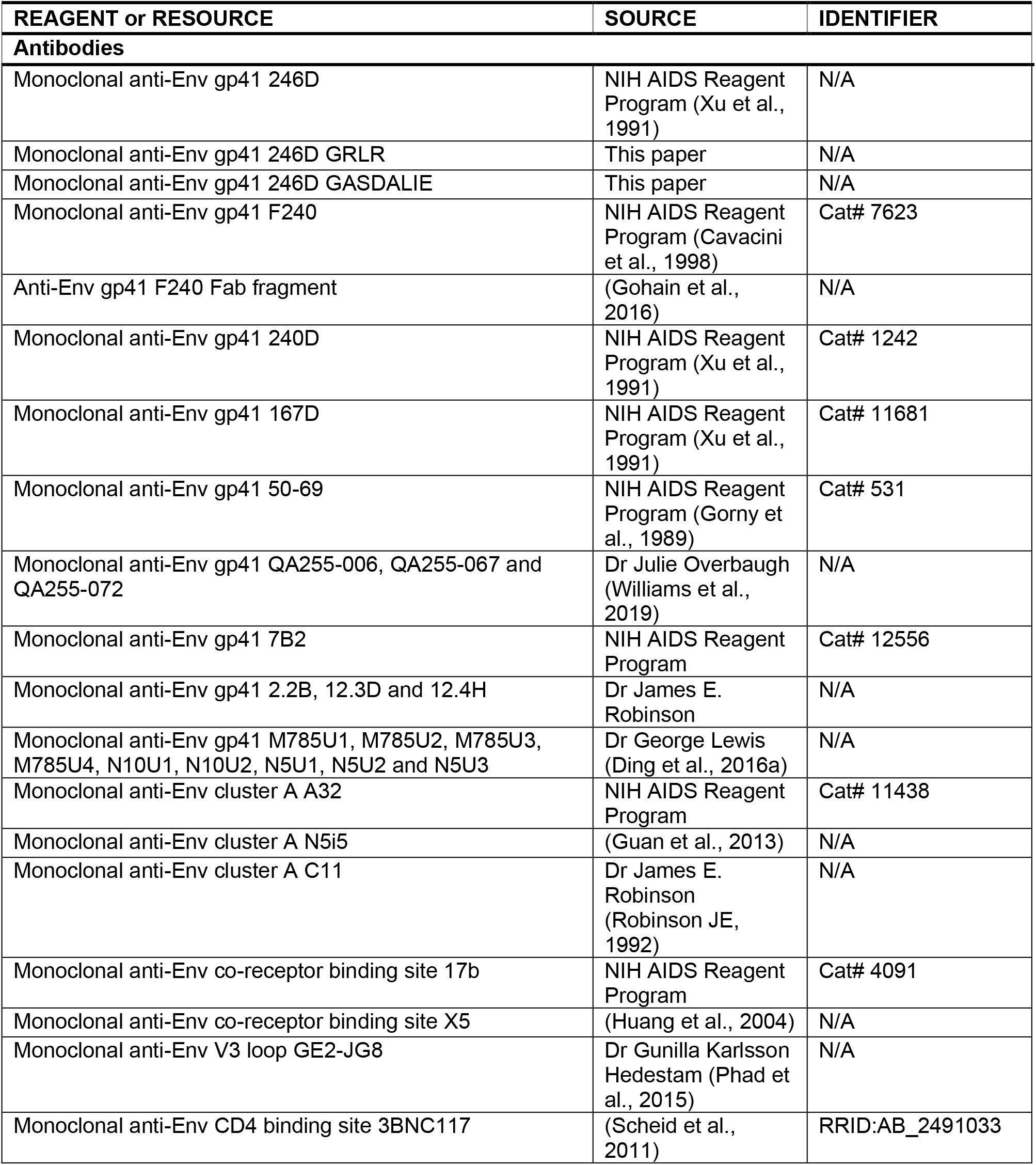
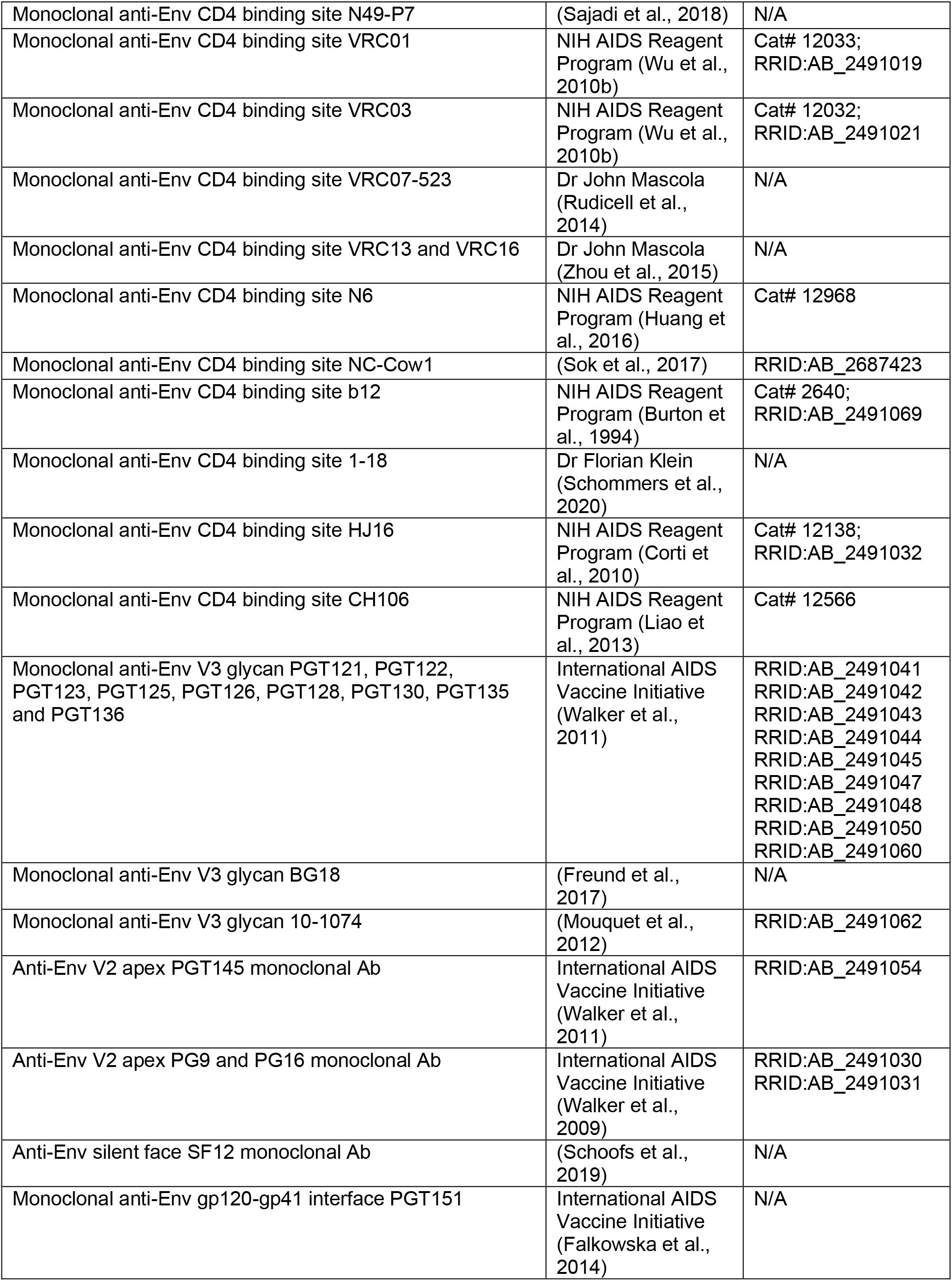
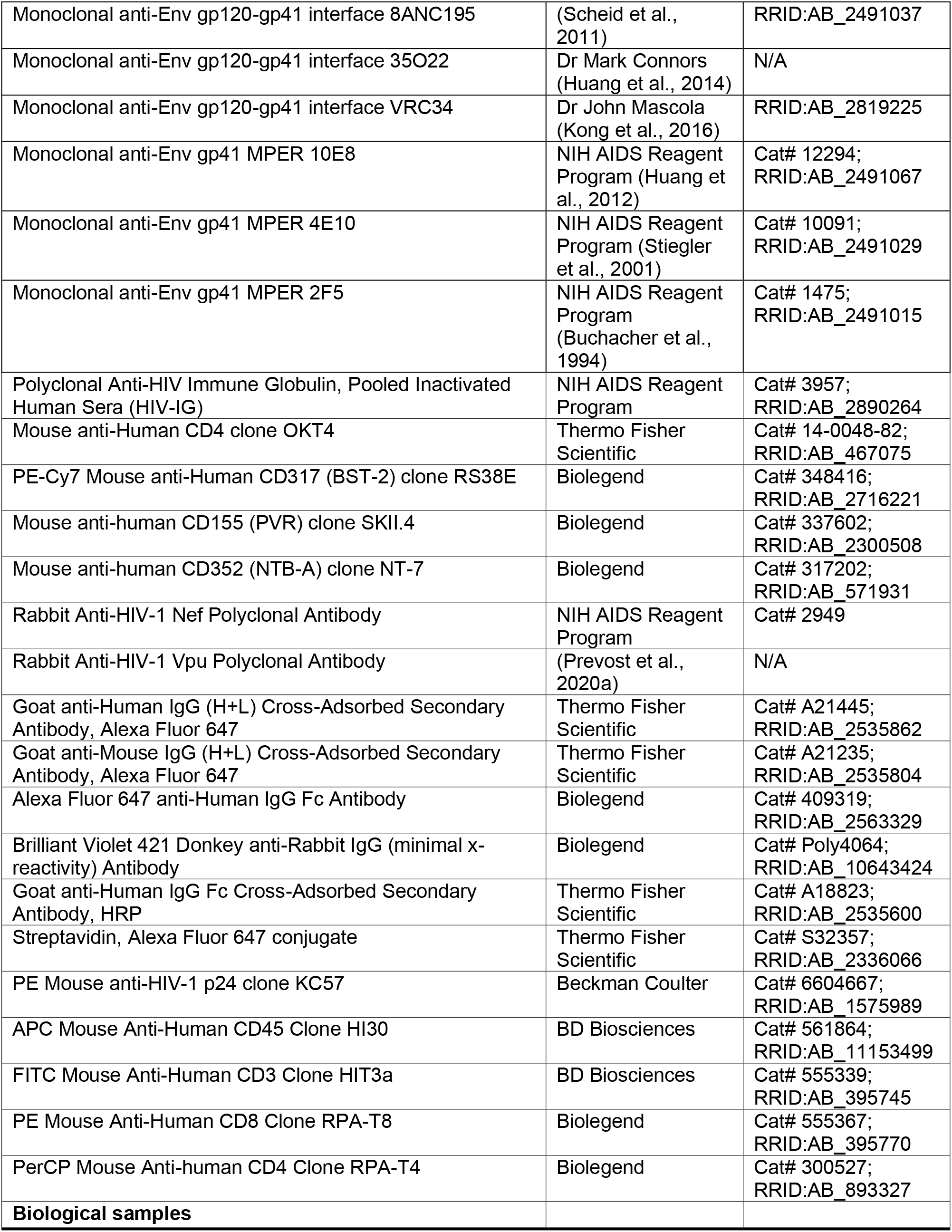
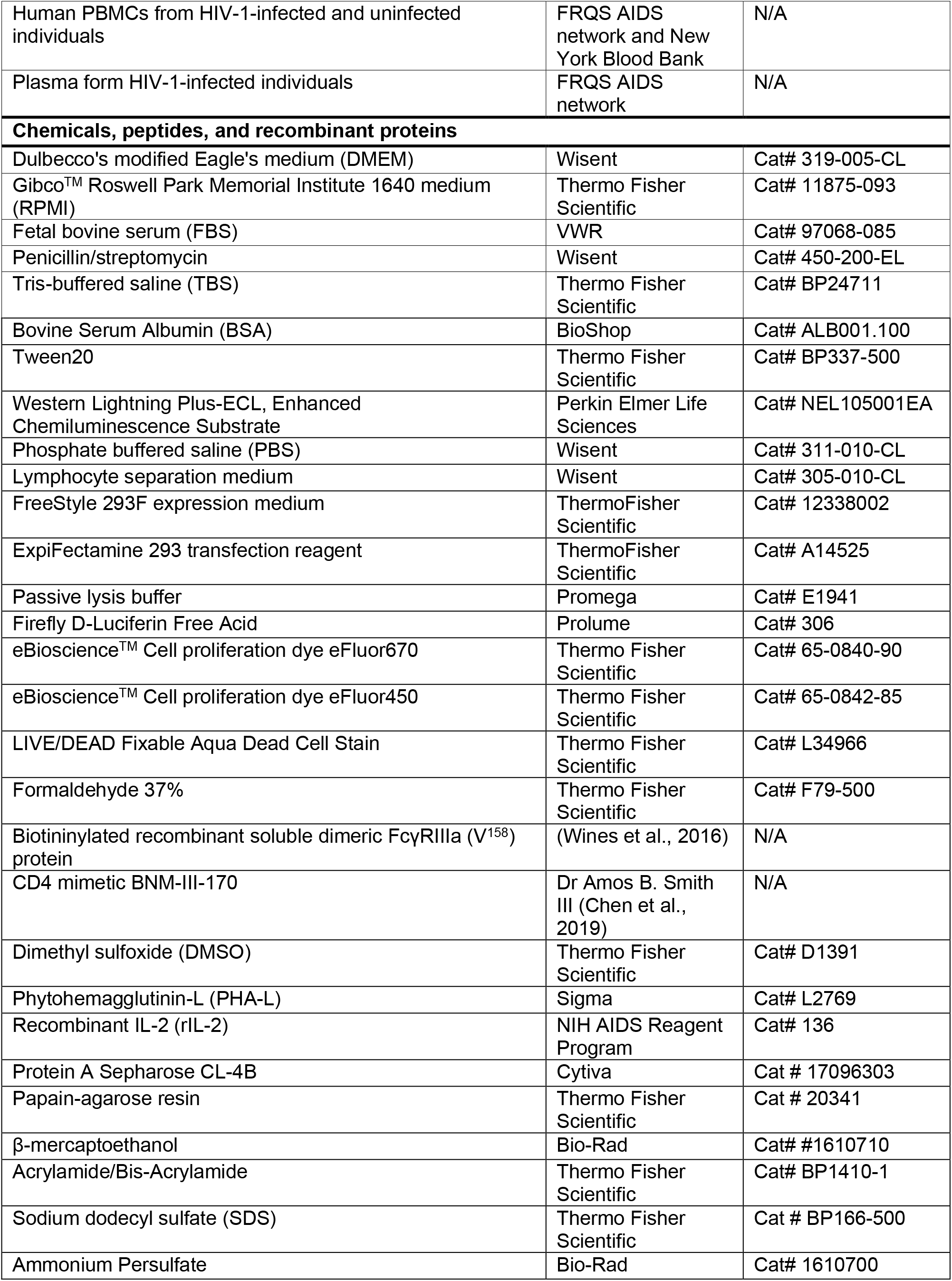
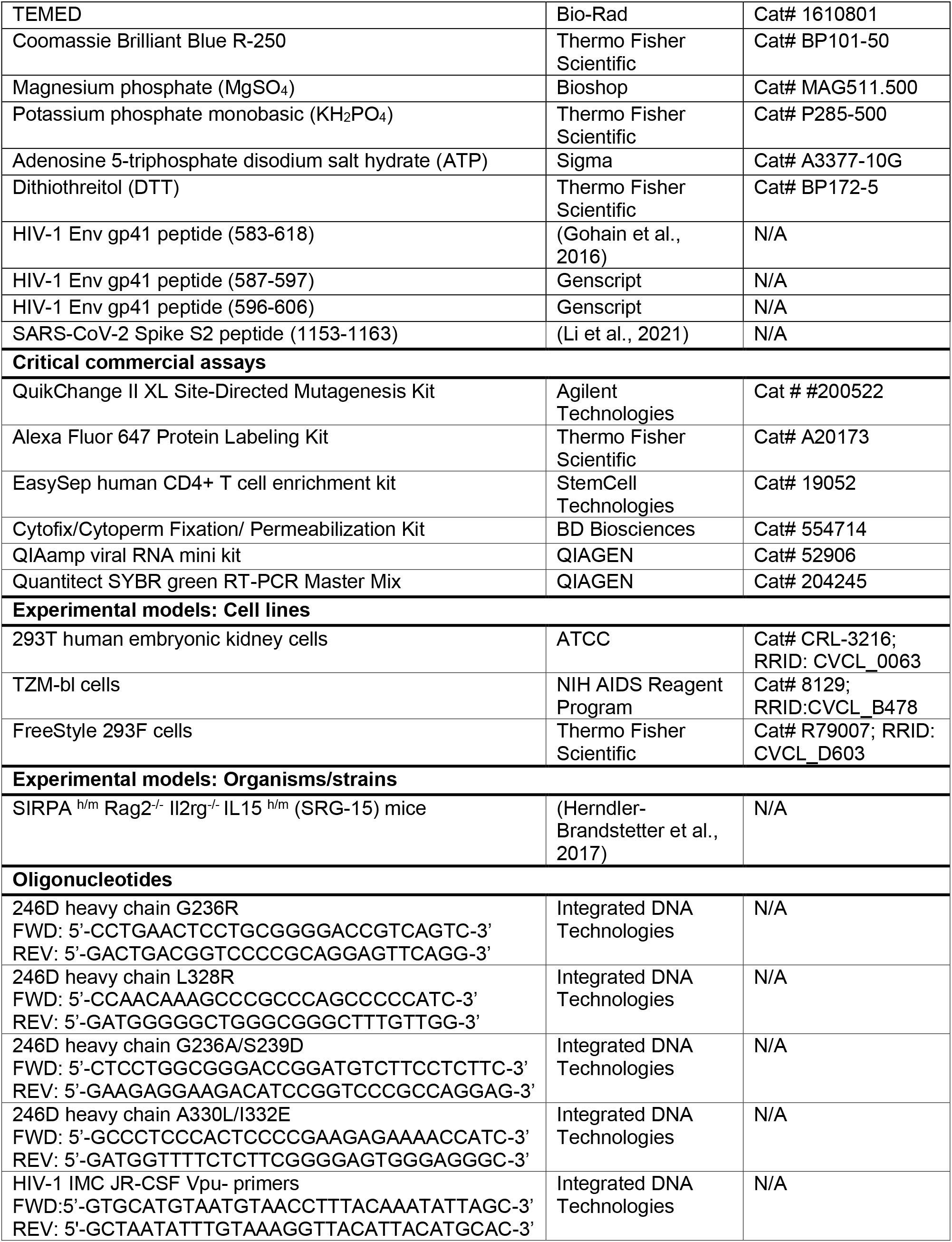
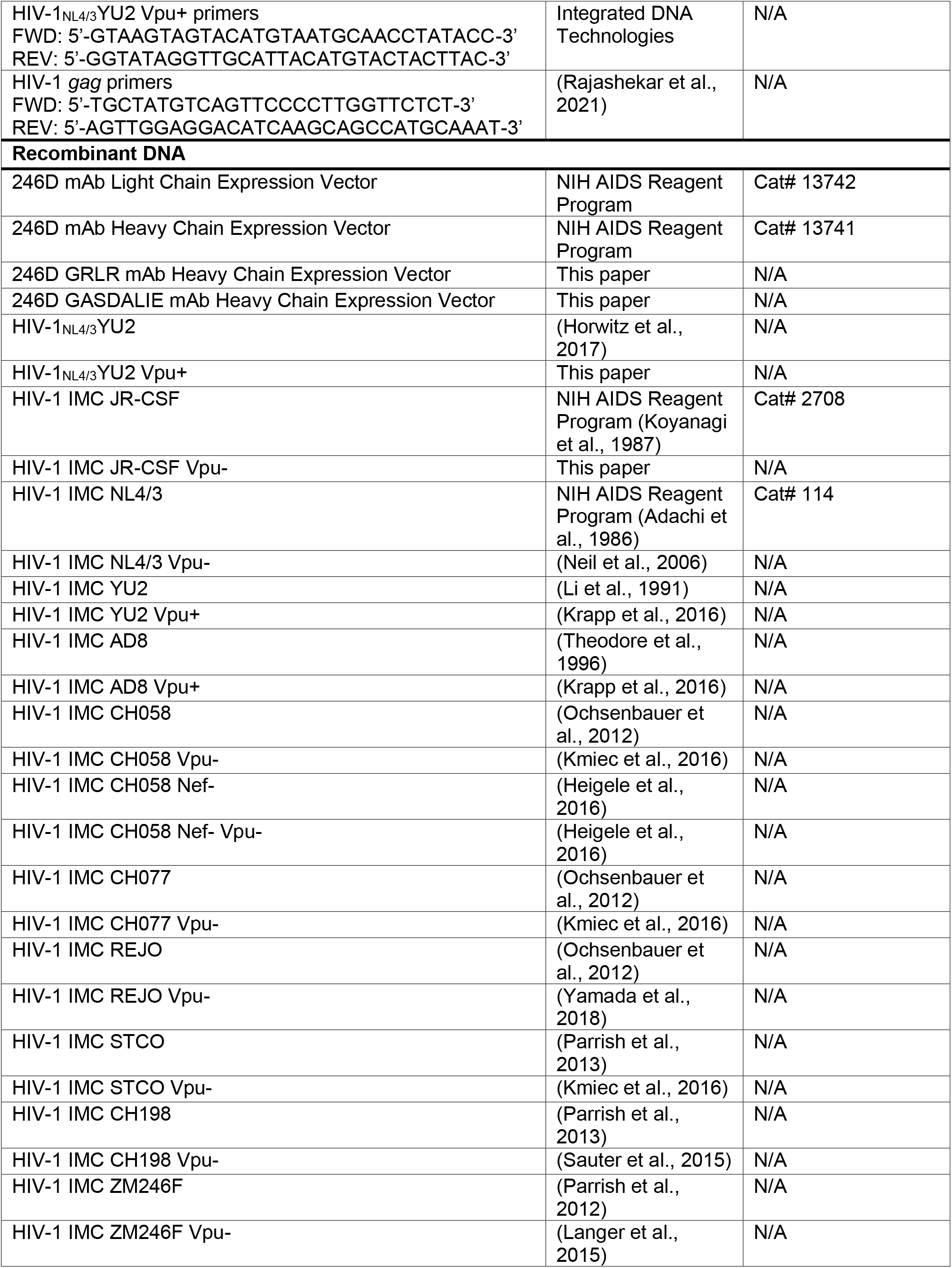
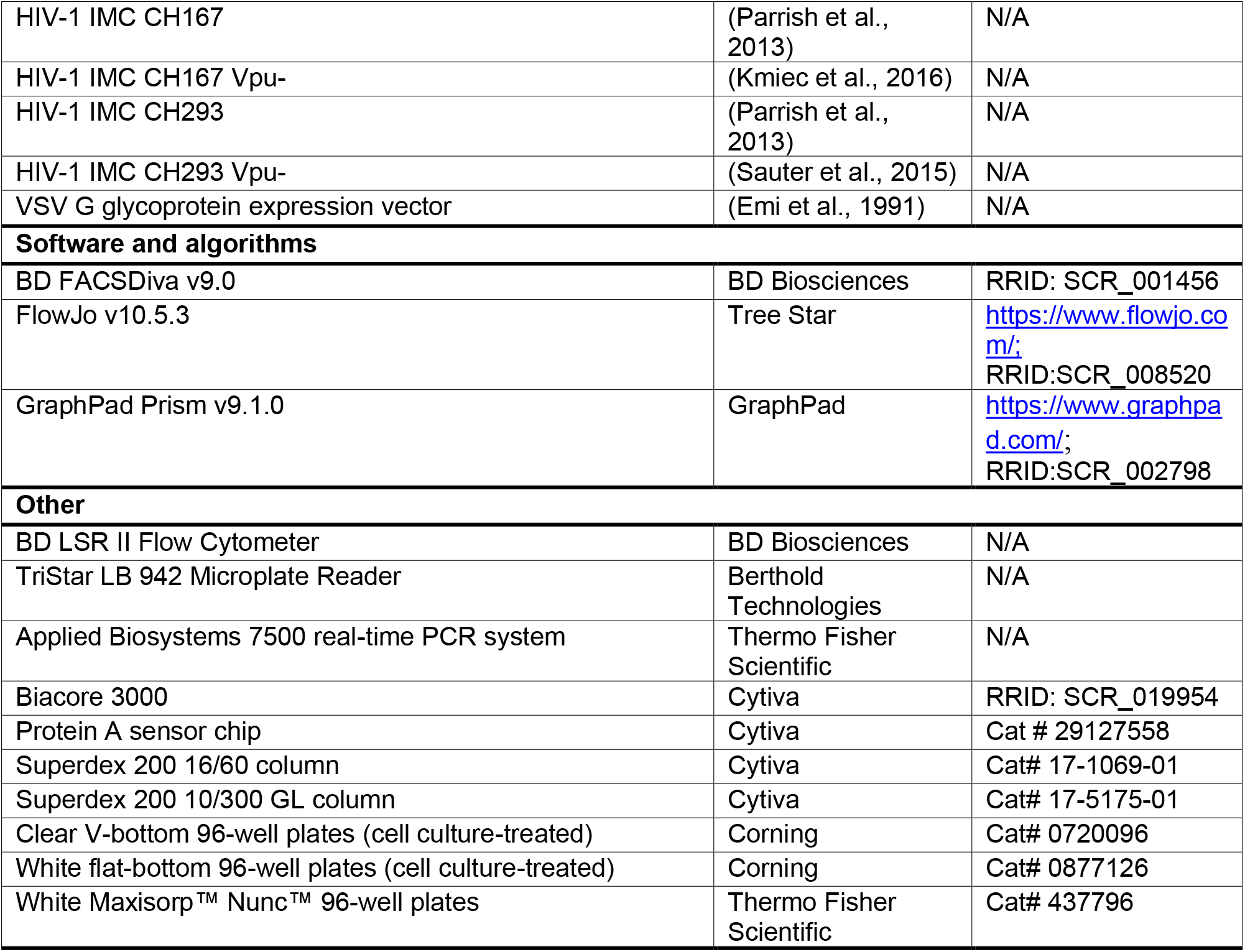

### RESOURCE AVAILABILITY

#### Lead Contact

Further information and requests for resources and reagents should be directed to and will be fulfilled by the Lead Contact, Andrés Finzi.

#### Materials Availability

All unique reagents generated in this study are available from the Lead Contact with a completed Materials Transfer Agreement.

#### Data and Code Availability

The published article includes all datasets generated and analyzed for this study. Further information and requests for resources and reagents should be directed to and will be fulfilled by the Lead Contact Author.

### EXPERIMENTAL MODELS AND SUBJECT DETAILS

#### Ethics Statement

Written informed consent was obtained from all study participants and research adhered to the ethical guidelines of CRCHUM and was reviewed and approved by the CRCHUM institutional review board (ethics committee, approval number CE 16.164 - CA). Research adhered to the standards indicated by the Declaration of Helsinki. All participants were adult and provided informed written consent prior to enrolment in accordance with Institutional Review Board approval.

#### Cell lines and primary cells

293T human embryonic kidney cells (obtained from ATCC) and TZM-bl cells (NIH AIDS Reagent Program) were maintained at 37°C under 5% CO_2_ in Dulbecco’s Modified Eagle Medium (DMEM) (Wisent, St. Bruno, QC, Canada), supplemented with 5% fetal bovine serum (FBS) (VWR, Radnor, PA, USA) and 100 U/mL penicillin/streptomycin (Wisent). 293T cells were derived from 293 cells, into which the simian virus 40 T-antigen was inserted. TZM-bl cells were derived from HeLa cells and were engineered to stably express high levels of human CD4 and CCR5 and to contain the firefly luciferase reporter gene under the control of the HIV-1 promoter (Platt et al., 1998). Human peripheral blood mononuclear cells (PBMCs) from ten HIV-negative individuals (8 males and 2 females) and five antiretroviral therapy (ART)-treated HIV-positive individuals (all males) obtained by leukapheresis and Ficoll-Paque density gradient isolation were cryopreserved in liquid nitrogen until further use. CD4+ T lymphocytes were purified from resting PBMCs by negative selection using immunomagnetic beads per the manufacturer’s instructions (StemCell Technologies, Vancouver, BC) and were activated with phytohemagglutinin-L (10 µg/mL) for 48 h and then maintained in RPMI 1640 (Thermo Fisher Scientific, Waltham, MA, USA) complete medium supplemented with rIL-2 (100 U/mL).

#### Human plasma samples

The FRQS-AIDS and Infectious Diseases Network supports a representative cohort of newly-HIV-infected subjects with clinical indication of primary infection [the Montreal Primary HIV Infection Cohort (Fontaine et al., 2011; Fontaine et al., 2009)]. Cross-sectional plasma samples from 50 HIV-1-infected individuals were segregated in five groups based on infection duration and treatment with antiretroviral therapy (ART). Plasma samples were obtained from treatment-naive subjects during the acute phase of infection (first 3 months after HIV acquisition), the early phase of infection (3–6 and 6-12 months after acquisition) and during the chronic phase of infection (>2 years after acquisition). Another group of chronically-infected individuals (>2 years after acquisition) received ART treatment. Plasma samples were also obtained from ten age- and sex-matched HIV-negative healthy volunteers.

#### Experimental animal models

SRG-15 mice encoding human SIRPA and IL-15 in a 129xBALB/c (N3) genetic background were originally generated in the laboratory of Dr Richard Flavell (Yale University) (Herndler-Brandstetter et al., 2017). The mice were bred and maintained under specific pathogen-free conditions. All animal studies were performed with authorization from Institutional Animal Care and Use Committees (IACUC) of Yale University. SRG-15-Hu-PBL mice were engrafted as described (Rajashekar et al., 2021). Briefly, 3.5 × 10^6^ PBMCs, purified by Ficoll density gradient centrifugation of healthy donor blood buffy coats (one male, obtained from the New York Blood Bank) were injected IP in a 200-µL volume into 6- to 8-week-old SRG-15 mice, using a 1-cm^3^ syringe and 25-gauge needle. Cell engraftment was tested 15 days post-transplant. 100 µL of blood was collected by retroorbital bleeding. PBMCs were isolated by Ficoll density gradient centrifugation; stained with fluorescently-labelled anti-human CD45, CD3, CD4, CD8 and CD56 antibodies and analyzed by flow cytometry to confirm engraftment. Humanized mice were intraperitoneally challenged with 30,000 PFU of HIV-1_NL4/3_YU2 Vpu- or Vpu+. Infection profile was analyzed through routinely by retro-orbital bleeds for plasma viral load (PVL) analysis.

### METHOD DETAILS

#### Plasmids and proviral constructs

The HIV-1_NL4/3_YU2 proviral construct has been previously reported (Horwitz et al., 2017). A mutation was introduced in the putative *vpu* start codon (ACG → ATG) to restore the *vpu* open reading frame (ORF) in the HIV-1_NL4/3_YU2 IMC using the QuikChange II XL site-directed mutagenesis protocol (Agilent Technologies, Santa Clara, CA). Transmitted/Founder (T/F) and chronic infectious molecular clones (IMCs) of patients CH058, CH077, CH198, ZM246F, CH167, CH293, REJO, STCO were inferred and constructed as previously described (Ochsenbauer et al., 2012; Parrish et al., 2013; Parrish et al., 2012; Salazar-Gonzalez et al., 2009). The generation of *vpu*-defective IMCs was previously described (Kmiec et al., 2016; Langer et al., 2015; Sauter et al., 2015; Yamada et al., 2018) and consists in the introduction of premature stop codons in the *vpu* reading frame using the QuikChange II XL site-directed mutagenesis protocol. CH058 IMCs defective for Vpu and/or Nef expression were previously described (Heigele et al., 2016; Kmiec et al., 2016). The IMCs encoding for HIV-1 reference strains NL4/3, AD8, YU2 and JR-CSF were described elsewhere (Adachi et al., 1986; Koyanagi et al., 1987; Krapp et al., 2016; Li et al., 1991; Theodore et al., 1996). To generate *vpu*-defective JR-CSF IMC, a stop-codon was introduced directly after the start-codon of *vpu* using the QuikChange II XL site-directed mutagenesis protocol. The plasmids encoding for the heavy and light chains of the 246D antibody are available through the NIH AIDS Reagent Program. Site-directed mutagenesis was performed on the plasmid expressing 246D antibody heavy chain to introduce the GRLR mutations (G236R/L328R) or the GASDALIE mutations (G236A/S239D/A330L/I332E) using the QuikChange II XL site-directed mutagenesis protocol. The presence of the desired mutations was determined by automated DNA sequencing. The vesicular stomatitis virus G (VSV-G)-encoding plasmid was previously described (Emi et al., 1991).

#### Viral production, infections and *ex vivo* amplification

For *in vitro* infection, vesicular stomatitis virus G (VSV-G)-pseudotyped HIV-1 viruses were produced by co-transfection of 293T cells with an HIV-1 proviral construct and a VSV-G-encoding vector using the calcium phosphate method. Two days post-transfection, cell supernatants were harvested, clarified by low-speed centrifugation (300 × g for 5 min), and concentrated by ultracentrifugation at 4°C (100,605 × g for 1 h) over a 20% sucrose cushion. Pellets were resuspended in fresh RPMI, and aliquots were stored at −80°C until use. Viruses were then used to infect activated primary CD4+ T cells from healthy HIV-1 negative donors by spin infection at 800 × *g* for 1 h in 96-well plates at 25 °C. Viral preparations were titrated directly on primary CD4+ T cells to achieve similar levels of infection among the different IMCs tested (around 10% of p24+ cells). To expand endogenously infected CD4+ T cells, primary CD4+ T cells obtained from ART-treated HIV-1-infected individuals were isolated from PBMCs by negative selection. Purified CD4+ T cells were activated with PHA-L at 10 μg/mL for 48 h and then cultured for at least 6 days in RPMI 1640 complete medium supplemented with rIL-2 (100 U/ml) to reach greater than 10% infection for the ADCC assay.

#### Antibodies

The following Abs were used to assess cell-surface Env staining: anti-gp41 nnAbs QA255-006, QA255-067, QA255-072 (kindly provided by Julie Overbaugh), 7B2, 2.2B, 12.3D, 12.4H (kindly provided by James Robinson), F240 (NIH AIDS Reagent Program), M785U1, M785U2, M785U3, M785U4, N10U1, N10U2, N5U1, N5U2, N5U3 (kindly provided by George Lewis), 246D, 240D, 167D, and 50-69; anti-cluster A A32 (NIH AIDS Reagent Program), C11 (kindly provided by James Robinson) and N5i5; anti-co-receptor binding site 17b (NIH AIDS Reagent Program) and X5; anti-V3 loop GE2-JG8 (kindly provided by Gunilla Karlsson Hedestam); anti-CD4 binding site VRC01, VRC03, VRC07-523, VRC13, VRC16 (kindly provided by John Mascola), N6 (kindly provided by Mark Connors), 1-18 (kindly provided by Florian Klein), HJ16, CH106 (NIH AIDS Reagent Program), NC-Cow1, b12, 3BNC117 and N49-P7; anti-V3 glycan PGT121, PGT122, PGT123, PGT125, PGT126, PGT128, PGT130, PGT135, PGT136 (IAVI), BG18 and 10-1074; anti-V2 apex PGT145, PG9 and PG16 (IAVI); anti-gp120-gp41 interface PGT151 (IAVI), 35O22 (kindly provided by Mark Connors), VRC34 (kindly provided by John Mascola) and 8ANC195; anti-silent face SF12; anti-MPER 10E8 (kindly provided by Mark Connors), 4E10 and 2F5 (NIH AIDS Reagent Program). The HIV-IG polyclonal antibody consists of anti-HIV immunoglobulins purified from a pool of plasma from HIV+ asymptomatic donors (NIH AIDS Reagent Program). Mouse anti-human CD4 (clone OKT4; Thermo Fisher Scientific), mouse anti-human BST-2 (clone RS38E, PE-Cy7-conjugated; Biolegend, San Diego, CA, USA), mouse anti-human NTB-A (clone NT-7, Biolegend) and mouse anti-PVR (clone SKII.4, Biolegend) were also used as primary antibodies for cell-surface staining. Goat anti-mouse IgG (H+L), goat anti-human IgG (H+L) (Thermo Fisher Scientific) and mouse anti-human IgG Fc (Biolegend) antibodies pre-coupled to Alexa Fluor 647 were used as secondary antibodies in flow cytometry experiments. Rabbit antisera raised against Nef (NIH AIDS Reagent Program) or Vpu (Prevost et al., 2020a) were used as primary antibodies in intracellular staining. BrillantViolet 421 (BV421)-conjugated donkey anti-rabbit antibodies (Biolegend) were used as secondary antibodies to detect Nef and Vpu antisera binding by flow cytometry. Goat anti-human IgG Fc antibodies conjugated to horseradish peroxidase (HRP; Thermo Fisher Scientific) were used as secondary antibodies in ELISA experiments.

#### Small CD4-mimetics

The small-molecule CD4-mimetic compound (CD4mc) BNM-III-170 was synthesized as described previously (Chen et al., 2019). The compounds were dissolved in dimethyl sulfoxide (DMSO) at a stock concentration of 10 mM and diluted to 50 µM in phosphate-buffered saline (PBS) for cell-surface staining or in RPMI-1640 complete medium for ADCC assays.

#### Protein production of recombinant proteins

FreeStyle 293F cells (Thermo Fisher Scientific) were grown in FreeStyle 293F medium (Thermo Fisher Scientific) to a density of 1 × 10^6^ cells/ml at 37 °C with 8 % CO_2_ with regular agitation (150 rpm). Cells were transfected with plasmids expressing the light and heavy chains of a given antibody using ExpiFectamine 293 transfection reagent, as directed by the manufacturer (Thermo Fisher Scientific). One week later, the cells were pelleted and discarded. The supernatants were filtered (0.22-μm-pore-size filter), and antibodies were purified by protein A affinity columns, as directed by the manufacturer (Cytiva, Marlborough, MA, USA). The recombinant protein preparations were dialyzed against phosphate-buffered saline (PBS) and stored in aliquots at −80°C. To assess purity, recombinant proteins were loaded on SDS-PAGE polyacrylamide gels in the presence or absence of β-mercaptoethanol and stained with Coomassie blue. Purified F240 and 2.2B IgG were conjugated with Alexa Fluor 647 dye according to the manufacturer’s protocol (Thermo Fisher Scientific). The F240 Fab fragments were prepared from purified IgG (10 mg/mL) by proteolytic digestion with immobilized papain (Pierce, Rockford, IL) and purified using protein A, followed by gel filtration chromatography on a Superdex 200 16/60 column (Cytiva). The biotin-tagged dimeric recombinant soluble FcγRIIIa (V^158^) protein was produced and characterized as described (Wines et al., 2016) with additional purification step using a Superdex 200 10/300 GL column (Cytiva).

#### Flow cytometry analysis of cell-surface staining

Cell surface staining was performed at 48h post-infection. Mock-infected or HIV-1-infected primary CD4+ T cells were incubated for 30 min at 37°C with anti-CD4 (0.5 µg/mL), anti-BST-2 (2 µg/mL), anti-NTB-A (5 µg/mL), anti-PVR (10 µg/mL), anti-Env mAbs (5 µg/mL), HIV-IG (50 µg/mL) or plasma (1:1000 dilution). Cells were then washed once with PBS and stained with the appropriate Alexa Fluor 647-conjugated secondary antibody (2 µg/mL), when needed, for 20 min at room temperature. After one more PBS wash, cells were fixed in a 2% PBS-formaldehyde solution. Alternatively, the binding of anti-Env Abs was detected using a biotin-tagged dimeric recombinant soluble FcγRIIIa (0.2 μg/mL) followed by the addition of Alexa Fluor 647-conjugated streptavidin (Thermo Fisher Scientific; 2 µg/mL). Anti-human IgG Fc secondary antibodies (3 µg/mL) were used when cell surface binding was performed in presence of F240 Fab blockade. Infected cells were then permeabilized using the Cytofix/Cytoperm Fixation/ Permeabilization Kit (BD Biosciences, Mississauga, ON, Canada) and stained intracellularly using PE-conjugated mouse anti-p24 mAb (clone KC57; Beckman Coulter, Brea, CA, USA; 1:100 dilution) in combination with Nef or Vpu rabbit antisera (1:1000 dilution). The percentage of infected cells (p24^+^) was determined by gating on the living cell population according to a viability dye staining (Aqua Vivid; Thermo Fisher Scientific). Alternatively, cells were stained intracellularly with rabbit antisera raised against Nef or Vpu (1:1000) followed by BV421-conjugated anti-rabbit secondary antibody. Samples were acquired on an LSR II cytometer (BD Biosciences), and data analysis was performed using FlowJo v10.5.3 (Tree Star, Ashland, OR, USA).

#### Antibody-dependant cellular cytotoxicity (ADCC) assay

Measurement of ADCC using a fluorescence-activated cell sorting (FACS)-based infected cell elimination (ICE) assay was performed at 48 h post-infection. Briefly, HIV-1-infected primary CD4+ T cells were stained with AquaVivid viability dye and cell proliferation dye eFluor670 (Thermo Fisher Scientific) and used as target cells. Cryopreserved autologous PBMC effectors cells, stained with cell proliferation dye eFluor450 (Thermo Fisher Scientific), were added at an effector: target ratio of 10:1 in 96-well V-bottom plates (Corning, Corning, NY). A 1:1000 final dilution of plasma or 5 µg/mL of anti-Env mAbs was added to appropriate wells and cells were incubated for 5 min at room temperature. The plates were subsequently centrifuged for 1 min at 300 × g, and incubated at 37 °C, 5 % CO_2_ for 5 h before being fixed in a 2 % PBS-formaldehyde solution. Infected cells were identified by intracellular p24 staining as described above. Samples were acquired on an LSR II cytometer (BD Biosciences) and data analysis was performed using FlowJo v10.5.3 (Tree Star). The percentage of ADCC was calculated with the following formula: [(% of p24+ cells in Targets plus Effectors) − (% of p24+ cells in Targets plus Effectors plus plasma) / (% of p24+ cells in Targets) × 100] by gating on infected lived target cells.

#### Virus neutralization assay

TZM-bl target cells were seeded at a density of 2 × 10^4^ cells/well in 96-well luminometer-compatible tissue culture plates (PerkinElmer, Waltham, MA, USA) 24 h before infection. HIV-1_NL4/3_YU2 Vpu- or Vpu+ (in a final volume of 50 µl) was pre-incubated with the indicated amounts of mAbs for 1 h at 37 °C before adding to the target cells. Following an incubation of 24 h at 37 °C, 100 µl of media was added to each well. Two days later, the medium was removed from each well, and the cells were lysed by the addition of 30 µl of passive lysis buffer (Promega, Madison, WI, USA) followed by one freeze-thaw cycle. An LB 942 TriStar luminometer (Berthold Technologies, Bad Wildbad, Germany) was used to measure the luciferase activity in each well after the addition of 100 µL of Luciferin buffer (15 mM MgSO_4_, 15 mM KH_2_PO_4_ [pH 7.8], 1 mM ATP, and 1 mM dithiothreitol) and 50 µL of 1 mM Firefly D-luciferin (Prolume, Pinetop, AZ, USA).

#### Plasma viral load measurements

Quantification of PVLs was done using the method described previously (Gibellini et al., 2004). Briefly, 100 µL of blood was collected from mice at each time point by retroorbital bleed. Plasma viral RNA was extracted using the QIAamp viral RNA mini kit (QIAGEN, Hilden Germany) following the manufacturer’s protocol. SYBR green real-time PCR assay was carried out in a 20-µL PCR mixture volume consisting of 10 µL of 2× Quantitect SYBR green RT-PCR Master Mix (QIAGEN) containing HotStarTaq DNA polymerase, 0.5 µL of 500 nM each oligonucleotide primer, 0.2 µL of 100× QuantiTect RT Mix (containing Omniscript and Sensiscript RTs), and 8 µL of RNA extracted from plasma samples or standard HIV-1 RNA (from 5 × 10^5^ to 5 copies per 1 mL). Highly conserved sequences on the gag region of HIV-1 were chosen, and specific HIV-1 gag primers were selected. The sequences of HIV-1 gag primers are 5’ TGCTATGTCAGTTCCCCTTGGTTCTCT 3’ and 5’AGTTGGAGGACATCAAGCAGCCATGCAAAT 3’. Amplification was done in an Applied Biosystems 7500 real-time PCR system, and it involved activation at 45 °C for 15 min and 95 °C for 15 min followed by 40 amplification cycles of 95 °C for 15 s, 60 °C for 15 s, and 72 °C for 30 s. For the detection and quantification of viral RNA, the real-time PCR of each sample was compared with threshold cycle value of a standard curve.

#### Sequence analysis

The LOGO plot (Crooks et al., 2004) was created using the Analyze Align tool at the Los Alamos National Laboratory - HIV database which is based on the WebLogo program (https://www.hiv.lanl.gov/content/sequence/ANALYZEALIGN/analyze_align.html) and the HIV-1 database global curated and filtered 2019 alignment, including 6,223 individual Env protein sequences. The relative height of each letter within individual stack represents the frequency of the indicated amino acid at that position. The numbering of Env amino acid sequences is based on the HXB2 reference strain of HIV-1, where 1 is the initial methionine.

#### gp41 peptide ELISA

Peptides corresponding to the gp41 C-C loop region (residues 583-618) were either previously described (Gohain et al., 2016) or purchased from Genscript (Piscataway, NJ, USA). A peptide corresponding to the SARS-CoV-2 Spike S2 stem helix (residues 1153-1163) was used as a negative control and was previously reported (Li et al., 2021). Peptides were prepared in PBS at a concentration of 1 μg/mL and were adsorbed to white 96-well plates (MaxiSorp Nunc) overnight at 4 °C. Coated wells were subsequently blocked with blocking buffer (Tris buffered saline [TBS] containing 0.1% Tween20 and 2% BSA) for 1 hour at room temperature. Wells were then washed four times with washing buffer (TBS containing 0.1% Tween20). Anti-gp41 246D and F240 or anti-gp120 A32 (50 ng/mL) were prepared in a diluted solution of blocking buffer (0.1 % BSA) and incubated with the peptide-coated wells for 90 minutes at room temperature. Plates were washed four times with washing buffer followed by incubation with horseradish peroxidase (HRP)-conjugated anti-human IgG secondary Ab (0.3 µg/mL in a diluted solution of blocking buffer [0.4% BSA]) for 1 h at room temperature, followed by four washes. HRP enzyme activity was determined after the addition of a 1:1 mix of Western Lightning Plus-ECL oxidizing and luminol reagents (Perkin Elmer Life Sciences). Light emission was measured with a LB942 TriStar luminometer (Berthold Technologies).

#### Surface plasmon resonance (SPR)

Surface plasmon resonance assays were performed on a Biacore 3000 (Cytiva) with a running buffer of 10 mM HEPES pH 7.5 and 150 mM NaCl, supplemented with 0.05% Tween20 at 25°C. The binding kinetics between the gp41 C-C loop and the 246D antibody were obtained in a format where 246D IgG was immobilized onto a Protein A sensor chip (Cytiva) with ∼300 response units (RU) and serial dilutions of gp41 (583-618) synthetic peptide were injected with concentrations ranging from 0.488 to 31.25 nM. The protein A chip was regenerated with a wash step of 0.1 M glycine pH 2.0 and reloaded with IgG after each cycle. Kinetic constants were determined using a 1:1 Langmuir model in bimolecular interaction analysis (BIA) evaluation software.

### QUANTIFICATION AND STATISTICAL ANALYSIS

Statistics were analyzed using GraphPad Prism version 9.1.0 (GraphPad, San Diego, CA, USA). Every data set was tested for statistical normality and this information was used to apply the appropriate (parametric or nonparametric) statistical test. P values <0.05 were considered significant; significance values are indicated as * P<0.05, ** P<0.01, *** P<0.001, **** P<0.0001.

## SUPPLEMENTAL FIGURE LEGENDS

**Table S1.** Cohort of HIV-1-infected individuals. Related to Figure 1.

| Group | Gender |  | Age (years)<sup>a</sup> | Days since infection<sup>a</sup> | Days since ART<sup>a</sup> | Days between infection and ART<sup>a</sup> | Viral load (copies/mL)<sup>a</sup> | CD4 count (cells/mm<sup>3</sup>)<sup>a</sup> |
| --- | --- | --- | --- | --- | --- | --- | --- | --- |
|  | Male | Female |  |  |  |  |  |  |
| Group 1<br>0-90 days untreated | 10 | 0 | 40<br>[18-55] | 68<br>[42-97] | N/A | N/A | 60848<br>[132-391113] | 470<br>[430-829] |
| Group 2<br>91-180 days untreated | 10 | 0 | 38<br>[20-58] | 135<br>[109-177] | N/A | N/A | 21663<br>[6641-195302] | 650<br>[420-1235] |
| Group 3<br>181-365 days untreated | 10 | 0 | 31<br>[24-45] | 240<br>[203-314] | N/A | N/A | 27785<br>[1866-260852] | 490<br>[230-770] |
| Group 4<br>2-5 years untreated | 11 | 1 | 36<br>[23-54] | 1158<br>[856-1810] | N/A | N/A | 29234<br>[14255-809600] | 421<br>[200-1311] |
| Group 5<br>2-5 years ART-treated | 7 | 1 | 37<br>[23-60] | 983<br>[794-1570] | 690<br>[19-879] | 428<br>[192-986] | 50<br>[40-3337] | 615<br>[170-1149] |

<sup>a</sup>Values shown are group medians with ranges in bracket.
ART: antiretroviral therapy
N/A: not applicable

**Figure S1.**
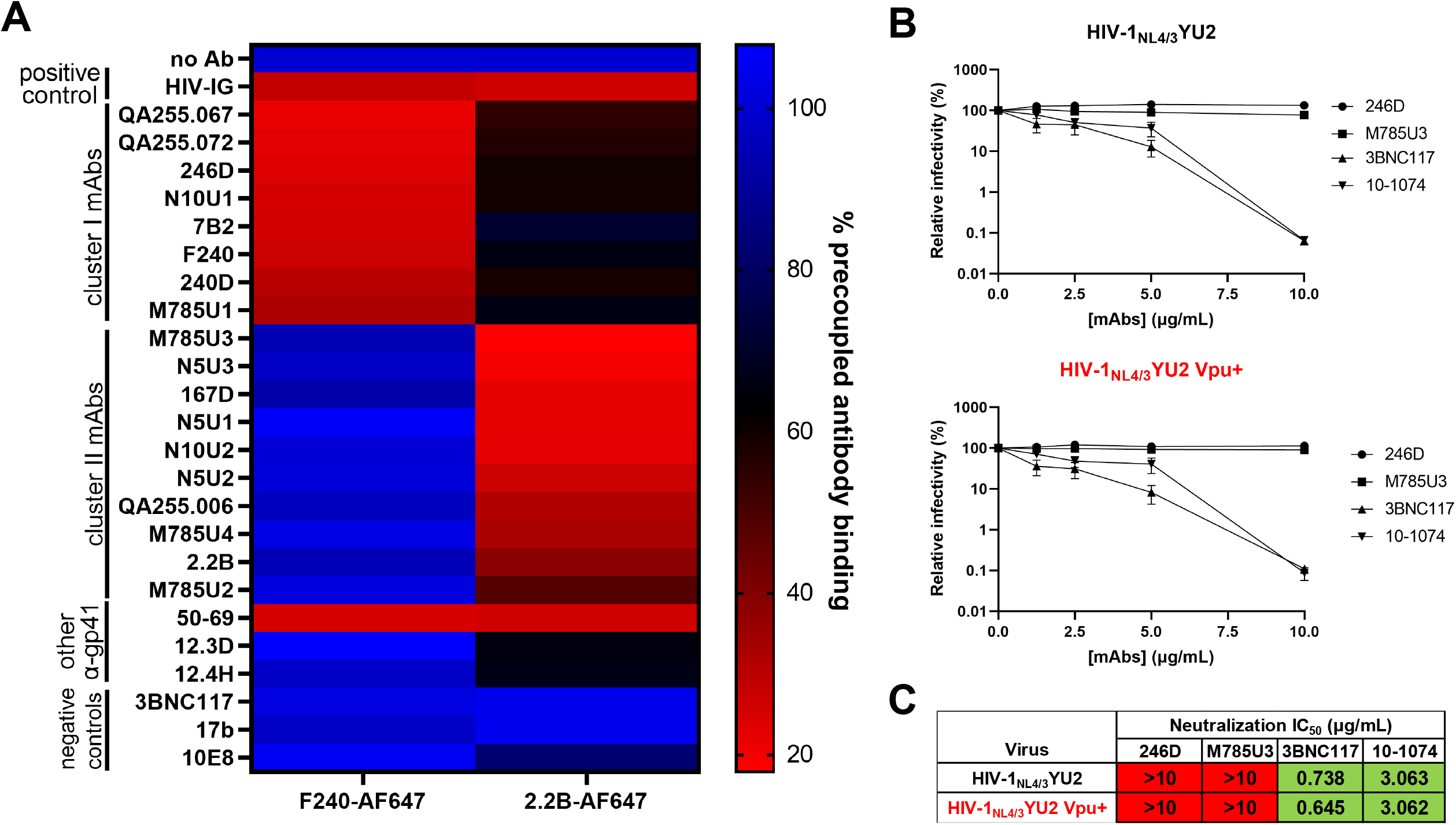
Classification of anti-gp41 non-neutralizing antibodies in two main clusters. Related to Figures 3 and 5. (**A**) The binding of Alexa Fluor 647 (AF647)-precoupled anti-gp41 cluster I F240 mAb or anti-gp41 cluster II 2.2B mAb was evaluated on HIV-1_NL4/3_YU2-infected cells in presence of a panel of unlabeled anti-gp41 nnAbs to determine epitope cross-competition. A pool of purified immunoglobulins from HIV-1-infected individuals (HIV-IG) was used as a positive control. Monoclonal antibodies with known non-competing epitopes (3BNC117, 17b, 10E8) were used as negative controls. (**B**) Lentiviral particles were produced from HIV-1_NL4/3_YU2 IMC expressing Vpu or not. Viruses were incubated with serial dilutions of anti-Env mAbs (246D, M785U3, 3BNC117, 10-1074) at 37°C for 1 h prior to infection of TZM-bl target cells. The infectivity at each Ab concentration tested is shown as the percentage of infection without Ab for each virus. Quadruplicate samples were analyzed in each experiment. The data shown are the means of results obtained in three independent experiments. Error bars indicate means ± the SEM. Neutralization half maximal inhibitory concentration (IC_50_) values are summarized in (**C**).

**Figure S2.**
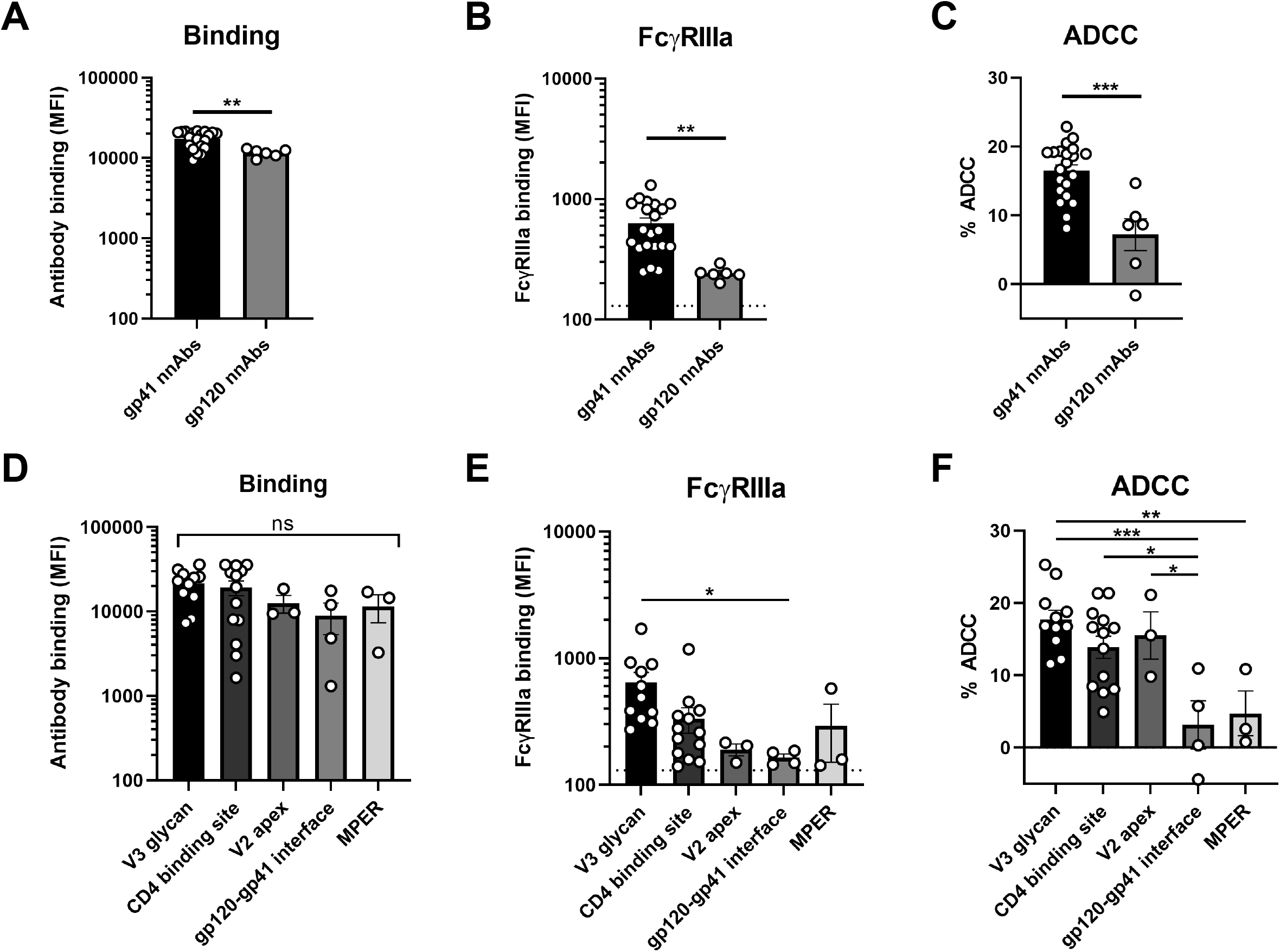
Epitope specificity dictates anti-Env ADCC responses mediated by nnAbs and bNAbs. Related to Figures 3 and 4. (**A-F**) Levels of antibody binding, FcγRIIIa binding and ADCC responses mediated by (**A-C**) nnAbs and (**D-F**) bNAbs as classified by epitope specificity (gp41 nnAbs, gp120 nnAbs; V3 glycan, CD4-binding site, V2 apex, gp120-gp41 interface, MPER). Statistical significance was tested using (**A-C**) an unpaired t test or a Mann-Whitney U test and (**D-F**) a one-way ANOVA with a Holm-Sidak post-test or a Kruskal-Wallis test with a Dunn’s post-test based on statistical normality (*, P < 0.05; **, P < 0.01; ***, P < 0.001; ****, P < 0.0001; ns, nonsignificant).

**Figure S3.**
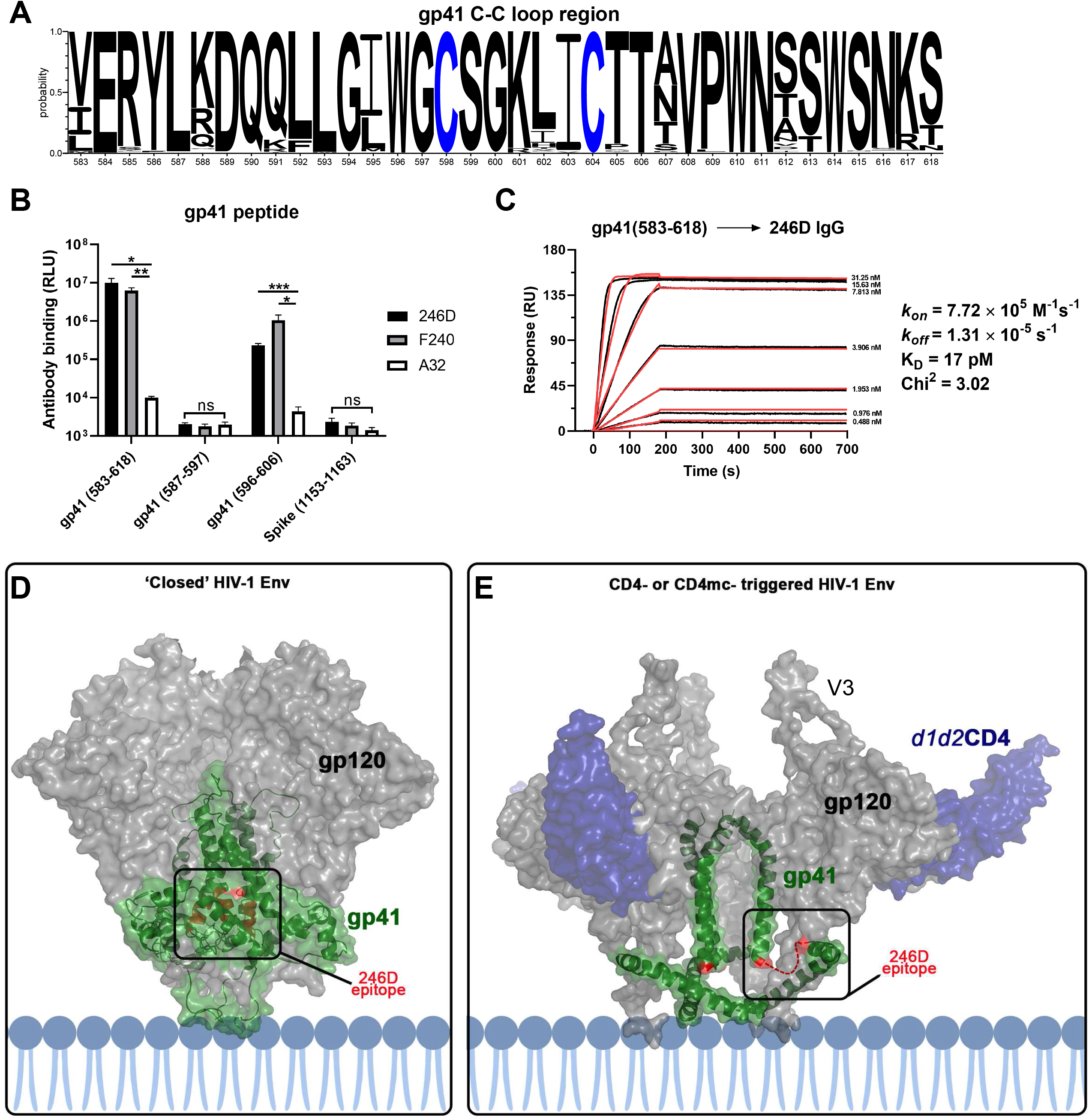
Monoclonal antibody 246D recognizes a gp41 linear peptide occluded in the closed Env trimer. Related to Figure 7. (**A**) Logo depiction of the frequency of each amino acid from the HIV-1 Env gp41 C-C loop region (residues 583-618) in all HIV-1 isolates. The height of the letter indicates its frequency among all strains. The 2019 Los Alamos database-curated filtered web Env alignment was used as the basis for this figure, which contains 6,223 individual Env amino acid sequences. Residue numbering is based on the HXB2 reference strain of HIV-1. (**B**) Indirect ELISA was performed using HIV-1 Env gp41 peptides corresponding to the C-C loop region, or a SARS-CoV-2 Spike S2 peptide as a negative control. Peptide-coated wells were incubated with anti-gp41 246D and F240 mAbs, as well as anti-gp120 cluster A A32 mAb as a negative control. Antibody binding was detected using HRP-conjugated anti-human IgG and was quantified by relative light units (RLU). The data shown are the means of results obtained in three independent experiments. Error bars indicate means ± the SEM. (**C**) 246D binding affinity and kinetics to gp41 C-C loop using surface plasmon resonance (SPR). The 246D IgG was immobilized as the ligand on a Protein A chip and HIV-1 gp41 (583-618) peptide used as analyte from 0.488 to 31.25 nM (2-fold serial dilution). Kinetic constants were determined using a 1:1 Langmuir model in bimolecular interaction analysis (BIA) evaluation software (experimental readings depicted in black and fitted curves in red). (**D-E**) Mapping of the ^596^WGCSGKLICTT^606^ epitope within available structures of HIV-1 Env. (**D**) The closed conformation of HIV-1 Env (PDB: 6ULC) (Pan et al., 2020) from a cryo-EM structure of full-length HIV-1 Env bound to the Fab of the antibody PG16 (not shown), with the 246D epitope highlighted in red. The 246D epitope is fully occluded in the closed the trimer and not accessible for antibody binding. (**E**) The CD4-triggered HIV-1 Env trimer (PDB: 3J70) (Rasheed et al., 2015) from a computational model of full-length HIV-1 Env bound to the d1d2 domain of CD4 and the Fab of antibody 17b (not shown). In the CD4 triggered trimer the 246D epitope is largely disordered (highlighted with a broken red line for one of three gp41 protomers), but it is exposed at the surface of trimer and available for antibody recognition.

